# The polyamine transporter ATP13A3 mediates DFMO-induced polyamine uptake in neuroblastoma

**DOI:** 10.1101/2024.02.20.581161

**Authors:** Mujahid Azfar, Weiman Gao, Chris Van den Haute, Lin Xiao, Mawar Karsa, Ruby Pandher, Emma Ronca, Angelika Bongers, Ayu Karsa, Dayna Spurling, Xinyi Guo, Chelsea Mayoh, Mark R. Burns, Steven H.L. Verhelst, Murray D. Norris, Michelle Haber, Peter Vangheluwe, Klaartje Somers

**Author notes:** these authors contributed equally (Mujahid Azfar, Weiman Gao). these authors contributed equally (Michelle Haber, Peter Vangheluwe, Klaartje Somers).

## Abstract

High-risk neuroblastomas, often associated with *MYCN* oncogene amplification, are addicted to polyamines, small polycations vital for cellular functioning. We have shown that neuroblastoma cells increase polyamine uptake when exposed to the polyamine biosynthesis inhibitor DFMO, currently in clinical trial, and that this mechanism limits the efficacy of the drug. While this finding resulted in the clinical development of polyamine transport inhibitors including AMXT 1501, presently under clinical investigation in combination with DFMO, the mechanisms and transporters involved in DFMO-induced polyamine uptake are unknown. Knockdown of ATP13A3, a member of the P5B-ATPase family, limited basal and DFMO-induced polyamine uptake, attenuated *MYCN*-amplified and non-*MYCN*-amplified neuroblastoma cell growth and potentiated the inhibitory effects of DFMO. Overexpression of ATP13A3 in neuroblastoma cells increased polyamine uptake, which was inhibited by AMXT 1501, highlighting ATP13A3 as a key target of the drug. The association between high ATP13A3 expression and poorer survival in neuroblastoma further supports a role of this transporter in neuroblastoma progression. Thus, this study identified ATP13A3 as a critical regulator of basal and DFMO-induced polyamine uptake and a novel therapeutic target for neuroblastoma.

**Graphical Abstract:** 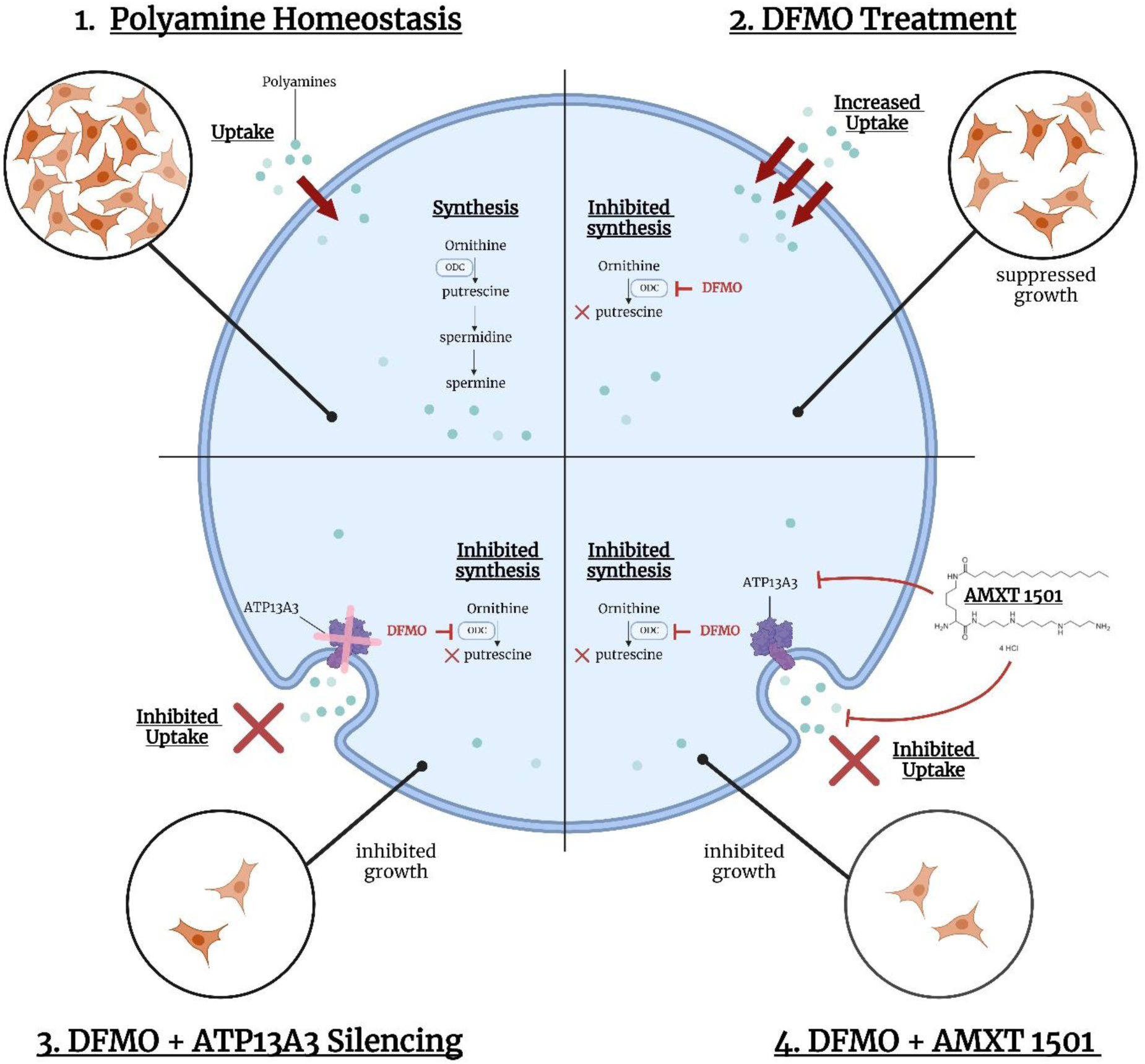

## 1. Introduction

Neuroblastoma, an embryonic cancer originating from neural crest cells and arising in the sympathetic nervous system, is the most common solid tumor among infants that causes around 15% of childhood cancer-related mortality ^1^ ^2^. High-risk neuroblastoma necessitates complex multi-modal treatments, including aggressive surgical resection, neoadjuvant chemotherapy, adjuvant high-dose chemotherapy with hematopoietic stem cell rescue, immunotherapy (anti-GD2 monoclonal antibody therapy) and radiation therapy ^3^. Despite this intensified treatment, a substantial number of high-risk patients do not respond well to conventional treatment, with more than half of children relapsing after therapy, the majority of whom ultimately die from the disease ^4^. Moreover, survivors of high-risk neuroblastoma often endure long-term treatment-related side effects, such as growth hormone deficiency, cataracts and reduced bone mineral density, drastically impairing their mental and physical quality of life^5^. Therefore, there is a pressing need for the development of safer and more precise therapeutic strategies, which in turn requires more insight into underlying disease mechanisms to identify better therapeutic targets.

One therapeutic strategy currently under investigation for cancer is targeting polyamine homeostasis. Polyamines, essential cations found in all living organisms, fulfil vital roles by interacting with a wide spectrum of negatively-charged molecules (*e.g.* RNA, DNA, phospholipids and proteins), thereby dictating the healthy functioning of cells ^6^. The three main polyamines found in mammalian cells are putrescine, spermidine and spermine. Polyamine homeostasis is heavily regulated by fine-tuning *de novo* synthesis, extracellular uptake, intracellular trafficking and catabolism. Because polyamines are regulators of cell growth, proliferation and survival, cancer cells rely on elevated polyamine levels to sustain their hyperproliferative state ^7^ ^8^. Accordingly, elevated polyamine levels are well-documented in cancer ^9^ ^7^ ^8^.

Neuroblastoma is also characterized by increased levels of intracellular polyamines, as well as elevated expression of ornithine decarboxylase 1 (ODC1), the rate limiting enzyme in the polyamine synthesis pathway, which is predictive of poorer outcome ^10^. In light of these findings, difluoromethylornithine (DFMO), a polyamine synthesis inhibitor targeting ODC1, and other agents targeting polyamine biosynthesis have been explored in pre-clinical neuroblastoma models, and were demonstrated to inhibit tumor growth ^11^. However, the first clinical investigations into DFMO in neuroblastoma yielded somewhat underwhelming results, and although DFMO maintenance therapy for high-risk neuroblastoma in remission appeared promising in reducing relapse and increasing overall survival time, the use of DFMO in the relapsed setting was not very effective ^12^ ^13^ ^14^. In exploring the disappointing results of DFMO in the relapsed setting, we found that neuroblastoma cells under a DFMO treatment regime compensate for reduced intracellular polyamine content by upregulating uptake of extracellular polyamines ^15^. We and others demonstrated that combination therapy that uses polyamine transport inhibitors such as AMXT 1501 in conjunction with DFMO, increases DFMO efficacy^16^ ^15^ ^17^. These results in pre-clinical models culminated in phase 1B/2A clinical trials for the combination of AMXT 1501 and DFMO (NCT03536728; NCT05500508). However, the molecular identity of the polyamine transporter(s) that is (are) targeted by AMXT 1501 and is (are) responsible for the DFMO-induced polyamine uptake remains unknown.

We previously uncovered a role for SLC3A2 in polyamine transport in neuroblastoma ^15^. We also established that members of the P5B-ATPase family (ATP13A2-4) are major polyamine transporters in the polyamine transport system ^18^ ^19^ that may be implicated in cancer ^6^ ^20^ ^21^, but their relative role in neuroblastoma remains unknown. In this study, we discovered that high ATP13A3 expression is associated with worse overall and event-free survival in neuroblastoma patients and that ATP13A3, but not SLC3A2, is a primary mediator of DFMO-induced polyamine uptake in this disease. Targeting ATP13A3, either by AMXT 1501 treatment or by silencing its expression, not only inhibits basal and DFMO-induced compensatory polyamine uptake and reduces the intracellular polyamine pool, but also abrogates colony formation capacity and growth of neuroblastoma cells while increasing the cells’ sensitivity to DFMO. Collectively our findings underpin ATP13A3 as not only a primary driver of DFMO-induced compensatory polyamine uptake in neuroblastoma but also a candidate target of AMXT 1501.

## 2. Material and Methods

### 2.1 Reagents

Polyamines putrescine dihydrochloride (P7505), spermidine (S2626) and spermine (3256) were purchased from SIGMA. All polyamines were dissolved to a final stock concentration of 500 mM (200 mM in the case of spermine) in 0.1 M MOPS-KOH (pH 7.0). AMXT 1501 was kindly provided by Aminex Therapeutics. Antibiotics (puromycin and blasticidin) for cell selection were purchased from INVIVOGEN.

### 2.2 Constructs and cell lines

SH-SY5Y cells were purchased from ATCC (CRL-2266, RRID:CVCL_0019), KELLY cells from DSMZ (ACC354, RRID:CVCL_2092), BE(2)-C cells (CRL-2268, RRID:CVCL_0529) from ATCC, SKNAS cells from ATCC (CRL-2137, RRID:CVCL_1700). All utilised cell lines were mycoplasma-free and have been authenticated in the past three years by STR profiling. To produce stable cell lines either overexpressing wildtype (WT) human ATP13A3 or a catalytically dead D498N (DN) mutant, cells were subjected to lentiviral transduction as previously described ^22^. Generation of stable ATP13A3 KD cell lines was also achieved using validated microRNA (miR) based short-hairpin lentiviral vectors targeting three different regions: miR2: AATCACAACAGATTCGTTATTT; miR3: TCAATCGTAAGCTCACTATATT; miR4: AGACCACCTTCGGGTCTTATAT. miRNA targeting Firefly Luciferase (mirFLUC: ACGCTGAGTACTTCGAAATGTC) was used as a negative control. Following transduction, the cells were kept under selection (2 µg/ml puromycin for overexpression models or 5 µg/ml blasticidin for the KD models). For siRNA knockdown of SLC3A2 and ATP13A3, four independent siRNAs were purchased from GE Dharmacon^TM^, and following optimization, the two or three siRNAS that gave the greatest knockdown were used for further experiments. siRNAs were transfected into cells using RNAiMAX according to manufacturer’s instructions. The parental SH-SY5Y cells, BE(2)-C cells and SK-N-AS cells were grown in 75 cm^2^ cell culture flasks in Dulbecco’s Modified Eagle’s Medium (DMEM) + 10% fetal calf serum (FCS) (Life Technologies), and stable ATP13A3 KD and overexpression SH-SY5Y cells were cultured in DMEM with high glucose supplemented with sodium bicarbonate, sodium pyruvate (SIGMA), Penicillin/streptomycin, L-Glutamine and 15% fetal bovine serum (PAN Biotech). KELLY cells were cultured in RPMI medium with 10-15% fetal calf serum (FCS) (Life Technologies). Cells were maintained at 5% CO_2_ and at 37°C and were split at full confluency (at 5% CO_2_ and at 37°C) using trypsin ethylenediaminetetraacetic acid solution (SIGMA) and phosphate-buffered saline (PBS) solution (SIGMA) without magnesium and calcium. The number of passages did not exceed 25.

### 2.3 Western Blotting

Total cell lysates were made in RIPA buffer (Thermo Fisher, 89900) containing protease inhibitors (Sigma, S8830). The protein concentration was determined using the Pierce BCA Protein Assay Kit (Thermo Fisher, 23227). Primary antibodies for ATP13A3 (Sigma, HPA029471, RRID: AB_10600784) and GAPDH (Sigma, G8795, RRID:AB_1078991) were dissolved at a dilution of 1:1000 and 1:5000 respectively, in 1% BSA/TBS-tween and incubated overnight at 4°C. Secondary antibodies, IgG HRP-linked Anti-rabbit (Cell Signaling, 7074S, RRID:AB_2099233) and IgG HRP-linked Anti-mouse (Cell Signaling, 7076S, RRID:AB_330924), were dissolved in 5% milk/TBS-tween solution at a dilution of 1:5000 for 1 hour. SuperSignal™West Pico PLUS chemiluminescent Substrate (Thermo Fisher, 34580) was used to detect bands, utilizing the Bio-Rad Chemidoc ™ MP imaging system.

### 2.4 PCR

RNA was isolated with RNeasy Micro kit (Qiagen, VIC, Australia). Quantitation of RNA was performed using NanoDrop™ 2000c Spectrophotometers (Thermo Scientific). Reverse transcription was performed with iScript™ Advanced cDNA Synthesis Kit (BioRad, NSW, Australia) and the PCR was performed with PrimePCR SYBR® Green Assays (BioRad) on a QuantStudio™ 3 Real-Time PCR System (Thermo Fisher Scientific, Australia). Forward primer sequence for ATP13A3: TACTGTGGAGCACTGATG; Reverse primer sequence for ATP13A3: GAGTTGCCACCATGTCATGC.

### 2.5 Cytotoxicity (viability) assays

The 4-methylumbelliferyl heptanoate (MUH) assay was performed as previously described ^18^. For MUH assays involving AMXT 1501, the cells were pre-treated overnight. Subsequent treatment with polyamines was performed in the presence of AMXT 1501.

### 2.6 Metabolomics

Intracellular polyamine levels were measured by the KU Leuven metabolomics core using LCMS. The samples were prepared using previously described protocols ^18^.

### 2.7 BODIPY-polyamine uptake assays

BODIPY-tagged polyamine probes were synthesized and provided by Prof. Dr. Steven Verhelst. Polyamine-BODIPY uptake was measured by flow cytometry as previously described^23^. For uptake assays involving AMXT 1501, the cells were pre-treated overnight. Subsequently, BODIPY-polyamines were added in the presence of AMXT 1501.

### 2.8 Radio-labelled polyamine uptake assays

^3^H-Spermidine Trihydrochloride (NET522001MC, 16.6 Ci/mmol) purchased from Perkin Elmer, and ^3^H-Putrescine Trihydrochloride (ART-0279-1mCi, 60 Ci/mol) obtained from American Radiolabeled Chemicals (via the Australian distributor Bio Scientific), were added to cells at 0.1-0.2 µCi, concentrations that obey the Michaelis Menten kinetics, and incubated at 37°C for 60-90 minutes prior to washing and lysis. Radiolabel incorporation was determined by scintillation counting and normalized to protein concentration as previously described ^15^.

### 2.9 Cell growth assays

Cells were seeded in a 12-well plate at 15k/well and transfected with siRNA at 24 hours post-seeding, followed by media replacement with or without drug treatment. Phase confluence cell images were obtained using a 10× objective lens every 6 hours for 168 hours and the average confluence of each well was calculated with the IncuCyte S3 Live Cell Analysis System (Sartorius, USA).

### 2.10 Colony formation assays

SH-SY5Y and KELLY cell lines were plated in 6-well plates at a density of 500-750 cells/well, and 24 hours after seeding were treated with a serial dilution of DFMO for 72 hours. After 10-14 days, cells were fixed and stained, and colonies counted using Image J software. Each experiment was performed in triplicate.

### 2.11 Synergy assays

For synergy viability assays, cells were seeded at 5-10k/well for 24 hours before being treated with increasing doses of drugs in a 6x6 combination matrix format. Cell viability was measured by resazurin reduction-based assays after a 3-day treatment. Synergy was scored according to the Bliss Independence model ^24^ and visualized by Combenefit ^25^.

### 2.12 ChIP-seq data processing and analysis

ChIP-seq fastq file data sets of MYCN in KELLY, BE(2)-C and NGP (RRID: RRID:CVCL_2141) cell lines (GSE80151) were obtained, as well as for c-MYC in NB69 cell line (GSE138295, RRID:CVCL_1448). Quality control analysis of sequencing reads was performed using fastQC (v0.11.9) and cutadapt (v4.2) was applied for adapter trimming. Reads were aligned against the GRCh38 human reference genome using bowtie2 (v2.5.0). BAM files were sorted and duplicates marked using picard (v2.27.5) and samtools (v1.12). Peak calling was performed using MACS2 (v 2.2.7.1), with narrow peak calling run with --keep-dup auto and -p 1e-9 for MYCN and -q 0.05 for c-MYC. ChIPseeker (v1.36.0) was employed for peak annotation with default parameters and the TxDb.Hsapiens.UCSC.hg38.knownGene transcript database. BigWig coverage files were generated using deeptools bamCoverage (v3.5.1).

### 2.13 Multivariate cox regression analysis

The Kocak microarray dataset (N=649) and SEQC RNA-seq dataset (N=498) were accessed through the R2: Genomics Analysis and Visualization Platform (R2: Genomics Analysis and Visualization Platform (amc.nl)). Event-free and overall survival data for the KOCAK dataset were supplied by the University of Cologne, Department of Pediatric Oncology and Hematology with clinical data such as patient age, tumor stage and *MYCN* amplification status retrieved from the Gene Expression Omnibus database (GSE45480). Kaplan-Meier survival curves were generated in R, and multi-variate analysis was performed with a cox proportional hazard model (using coxph function in R). *ATP13A3* expression was separated into either high (top 25th percentile) or low (bottom 75th percentile) and patient’s age into either ≥18 years or <18 years. The SEQC dataset covariates were *ATP13A3* expression (high versus low), age (≥18 versus <18 months), and *MYCN* status (amplified versus non-amplified). Kocak covariates also included tumor stage (stage 3/4 versus stage 1/2/4S).

### 2.14 Statistics

Data is presented as mean ± SEM of three independent repeats (unless otherwise stated). All statistical analyses were performed with GraphPad Prism. Statistical significance was determined by one-or two-way ANOVA followed by multiple comparisons using Dunnett’s/Tukey’s *post hoc* tests. For datasets comparing two groups, unpaired t-tests or one sample t-tests were performed. Statistical significance was defined as * p < 0.05, ** p < 0.01, *** p < 0.001, ****, p<0.0001 or ns (non-significant).

### 2.15 Data Availability

The data generated in this study are available upon request from the corresponding authors.

## 3. Results

### 3.1. SLC3A2-mediated polyamine transport does not contribute to DFMO-induced compensatory polyamine uptake

We previously identified SLC3A2 as a candidate polyamine transporter in neuroblastoma cells ^15^. Survival analyses in a neuroblastoma patient cohort (SEQC, n=498) confirmed our prior findings that elevated expression of *SLC3A2* is associated with worse event-free and overall survival **(SF 1)**. We also confirmed and extended our earlier discovery that SLC3A2 contributes to polyamine uptake in *MYCN*-amplified neuroblastoma cells ^15^, since siRNA-mediated *SLC3A2* silencing significantly reduced putrescine and spermidine uptake in non-*MYCN*-amplified (SH-SY5Y) as well as *MYCN*-amplified (KELLY) neuroblastoma cells **(Figure 1A – B, SF2)**. In light of the finding that neuroblastoma cells treated with DFMO compensate for a decrease in intracellular polyamines by augmenting polyamine uptake, we next assessed whether SLC3A2-mediated polyamine uptake is responsible for the DFMO-induced increase in polyamine uptake ^15^ ^26^ ^27^. We first confirmed that DFMO treatment significantly increased uptake of radiolabeled putrescine and spermidine in non-*MYCN* amplified and *MYCN*-amplified neuroblastoma cells (**Figure 1A – B**). Knockdown of SLC3A2 did not abrogate the DFMO-induced increase in polyamine uptake in *MYCN*-amplified (KELLY) or non-*MYCN*-amplified (SH-SY5Y) cells **(Figure 1A – B),** indicating that SLC3A2 alone does not drive the compensatory increase in polyamine uptake caused by DFMO. This finding provided impetus to investigate other polyamine transporters as potential contributors to polyamine transport in neuroblastoma and to polyamine uptake mechanisms that limit responsiveness to DFMO.

**Fig 1:**
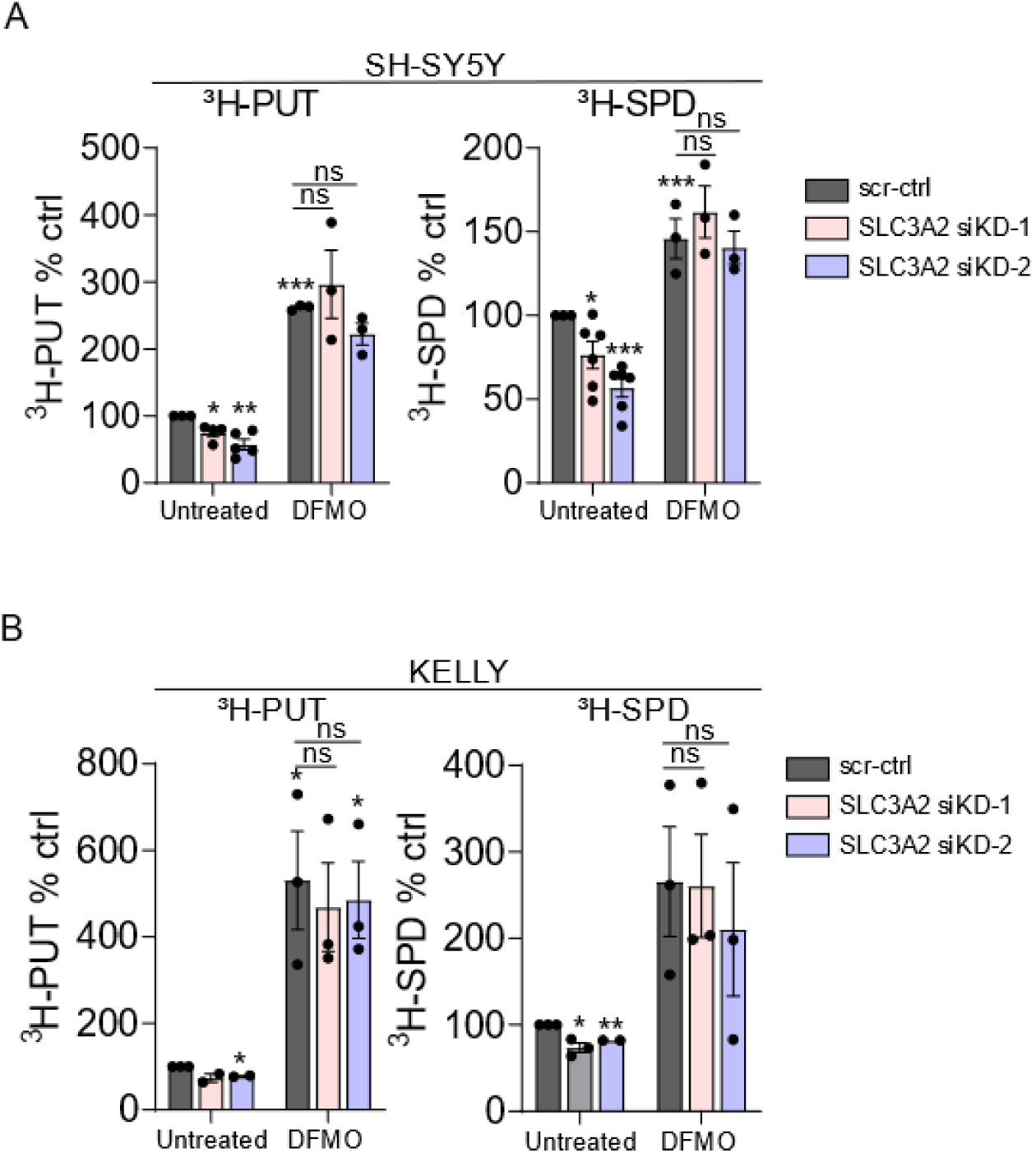
SLC3A2 silencing does not abrogate DFMO-induced polyamine uptake in neuroblastoma cells. (A – B) Measurement of radiolabelled putrescine (PUT) or spermidine (SPD) uptake after SLC3A2 silencing with two different siRNAs (siKD-1, siKD-2) versus a scrambled control siRNA (scr-ctrl). One sample t-test was used to assess the significance of the changes in polyamine uptake vs. untreated cells transfected with scr-ctrl (100%) which is indicated by the stars above the bars. Comparisons between the uptake % in all DFMO-treated groups were made by one-way ANOVA. Graphs depict mean +/- SEM of at least three independent biological replicates.

### 3.2. High *ATP13A3* expression is a prognostic predictor of poor outcome in neuroblastoma

We next explored a new class of transporters, the P5B-ATPase family, that we have recently established as key members of the mammalian polyamine transport system ^18^ ^19^ ^21^. To evaluate a role for P5B-ATPases in neuroblastoma, we first assessed their expression in neuroblastoma patients using the R2: Genomics Analysis and Visualization Platform (https://r2.amc.nl). Of the P5B-ATPases, only *ATP13A2* and *ATP13A3* are abundantly expressed in neuroblastoma tumours, similar to *SLC3A2* **(Figure 2A)**. Elevated *ATP13A3* but not *ATP13A2* expression was associated with worse event-free and overall survival in neuroblastoma as a whole, as well as in the *MYCN*-amplified neuroblastoma subpopulation **(Figure 2B-C, SF 3, SF 4)**, thus ATP13A3 was chosen for further evaluation.

**Figure 2:**
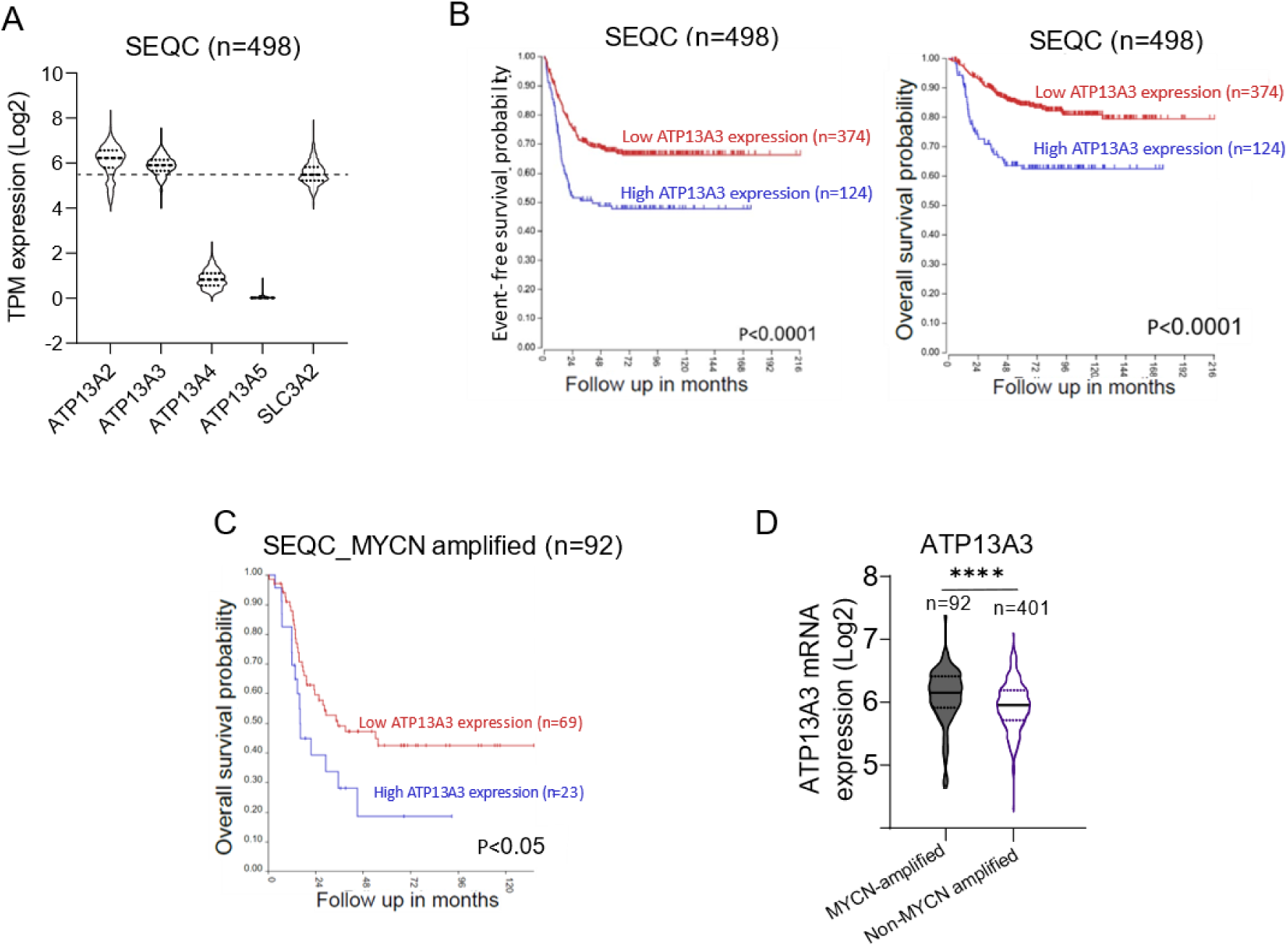
High expression of *ATP13A3* is a prognostic predictor for poor outcome in neuroblastoma. **(A)** mRNA expression (Transcripts Per Million, TPM) of polyamine transporters in the SEQC neuroblastoma database (n=498). **(B)** High expression of *ATP13A3* is associated with worse prognosis in the entire neuroblastoma cohort (SEQC database n=498) and **(C)** within the *MYCN*-amplified neuroblastoma patient subset (n=92). Patients were dichotomized around the upper quartile (UQ) of gene expression. Log-rank test was used to compare the survival curves of the high and low expression groups. **(D)** *ATP13A3* expression is higher in neuroblastoma patients with *MYCN* amplification. Student’s t-test was used for statistical analysis.

Interestingly, *ATP13A3* expression was significantly higher in *MYCN*-amplified versus non-*MYCN* amplified patients **(Figure 2D).** As *MYCN* gene amplification is the strongest genetic predictor for poor prognosis in neuroblastoma and has been shown to drive polyamine synthesis and uptake with *ODC1* and *SLC3A2* being identified as *bona fide* MYCN targets ^31^ ^32^ ^15^, we assessed whether the observed association between high *ATP13A3* expression and poor outcome is in fact mediated by *MYCN* amplification. Multivariate survival analyses on two neuroblastoma cohorts (Kocak GSE45547; SEQC GSE62564) which incorporated *MYCN* amplification, patient age and tumor stage as other prognostic variables, indicated that *ATP13A3* is an independent prognostic marker for neuroblastoma outcome (**Table 1**). We next assessed whether *ATP13A3* is a target gene of the family of MYC transcription factors, similar to *ODC1* and *SLC3A2*. We analyzed publicly available ChIP-Seq databases and found that both MYCN and c-MYC bind to the promotor of *ATP13A3*, suggesting that *ATP13A3* is a MYC target gene similar to *ODC1* and *SLC3A2* **(SF 5 and SF 6)**.

**Table 1:**
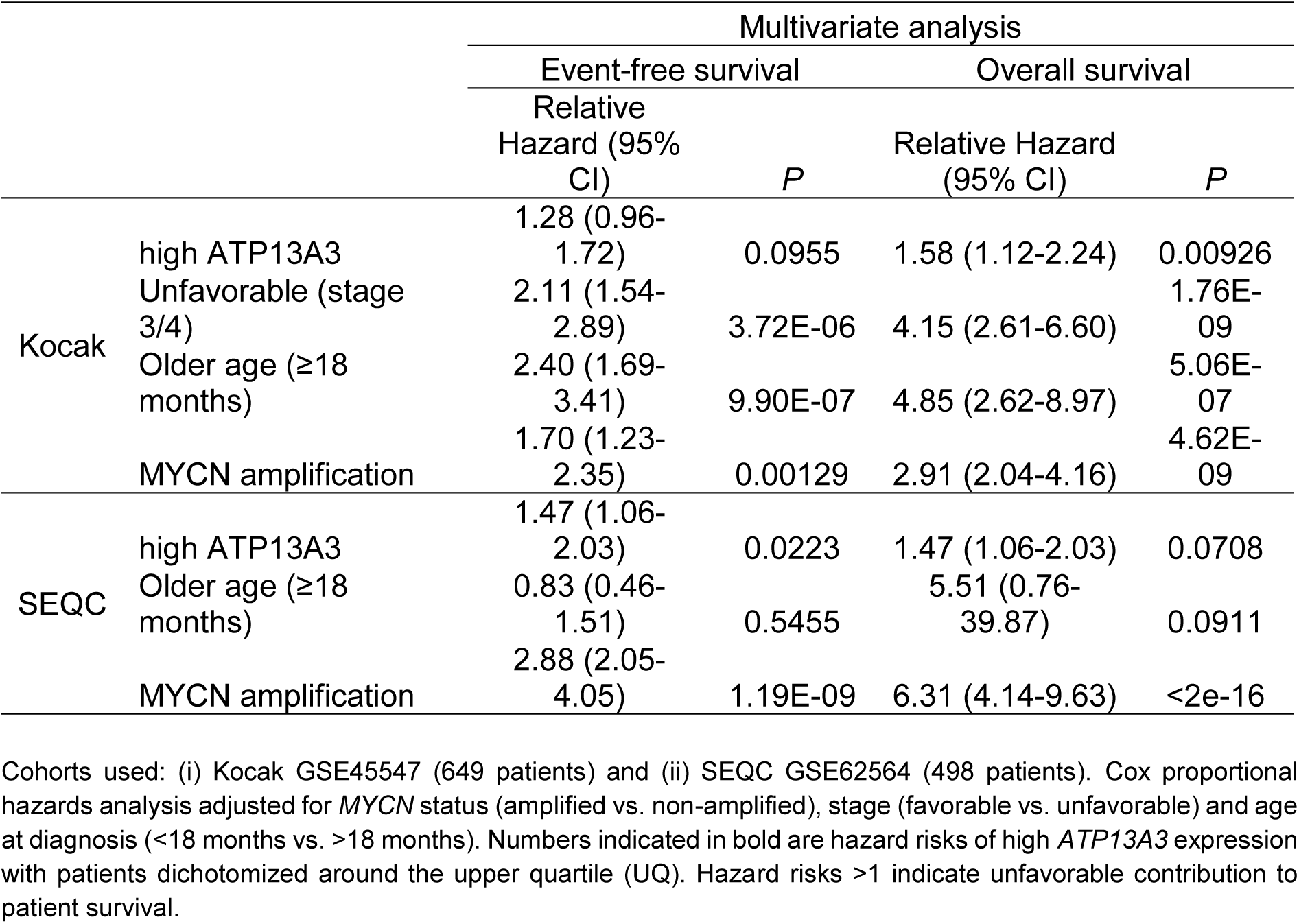
Multivariate analysis for high ATP13A3 expression showing the event-free and overall survival data in two neuroblastoma patient cohorts.

Our findings thus indicate that high expression of *ATP13A3* represents an independent prognostic predictor of poor outcome in neuroblastoma, suggesting a role for ATP13A3 in neuroblastoma progression, and identify *ATP13A3* as a candidate MYC target gene.

### 3.3 ATP13A3 mediates polyamine transport in neuroblastoma cells irrespective of MYCN status

To study the role of ATP13A3 in polyamine uptake and homeostasis, we generated SH-SY5Y cells that overexpress wildtype (WT) ATP13A3 and ATP13A3 D498N (DN) – a catalytically dead/transport deficient mutant in which the catalytic site for reversible phosphorylation (Asp498) is replaced by asparagine **(SF 7)**. We observed that cells that overexpressed WT ATP13A3 contained higher basal intracellular levels of all three polyamines **(Figure 3A)**. To demonstrate that these higher polyamine levels were due to increased ATP13A3-mediated polyamine uptake ^19^ ^20^, we assessed the uptake of fluorescent BODIPY-conjugated polyamines, that behave similarly to radiolabeled polyamines ^23^. We noted a significant increase in putrescine-BODIPY (PUT-BDP) and spermidine-BODIPY (SPD-BDP), but not in spermine-BODIPY (SPM-BDP) uptake upon WT ATP13A3 overexpression compared to DN ATP13A3 overexpression **(Figure 3B)**.

**Figure 3:**
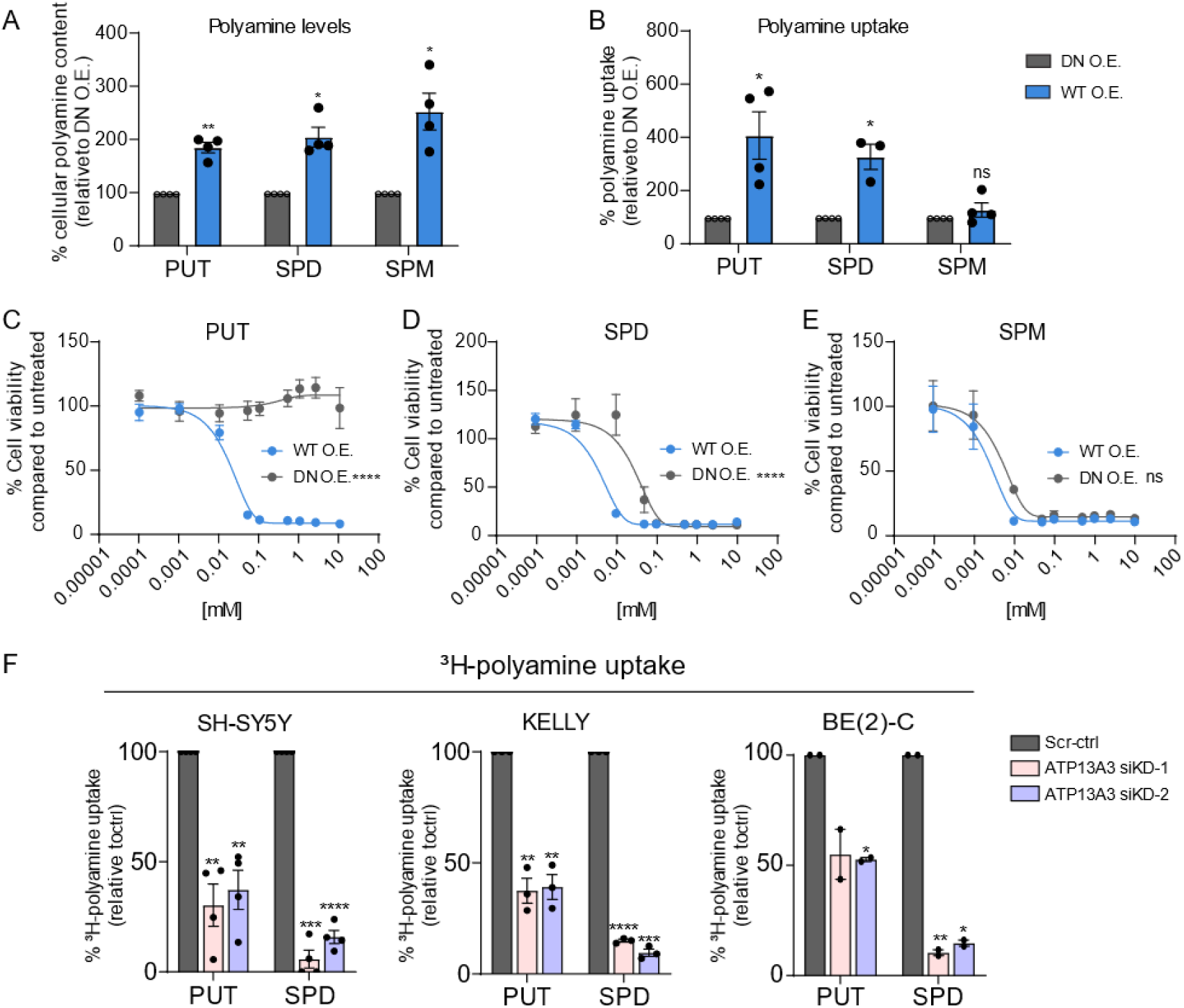
ATP13A3 mediates polyamine uptake and homeostasis in neuroblastoma cells. **(A)** Mass spectrometry-based measurement of polyamine levels demonstrating that the overexpression (O.E.) of WT ATP13A3 in SH-SY5Y cells increases the basal level of putrescine (PUT), spermidine (SPD) and spermine (SPM) when compared to DN overexpression. One sample t-test was used for statistical analysis. Graphs depict mean +/- SEM of at least four independent biological replicates. **(B)** Overexpression of WT ATP13A3 stimulates the uptake of fluorescent BODIPY (BDP)-conjugated PUT and SPD but not of SPM in SH-SY5Y cells compared to cells overexpressing DN ATP13A3. One sample t-test was used to determine the significance of the difference in uptake. Graphs depict mean +/-SEM of at least four independent biological replicates. **(C – E)** Cell viability assay, using the MUH reagent to assess, showing that overexpression of WT ATP13A3 sensitizes SH-SY5Y cells to toxic effects of exogenously added PUT and SPD when compared to DN overexpression. No significant difference was seen in SPM-conferred toxicity between WT and DN overexpressing cells. Two-way ANOVA was used for statistical analysis. Graphs depict mean +/-SEM of at least two independent biological replicates (PUT: n=3; SPD: n=3; SPM: n=2). (F) ATP13A3 siRNA-mediated knockdown (KD) reduces ^3^H-labelled PUT and SPD uptake in non-*MYCN* amplified (SH-SY5Y) and *MYCN*-amplified (KELLY and BE(2)-C) neuroblastoma cells. One sample t-test was used to assess the significance of the decrease in uptake compared with Scr-ctrl cells. Graphs depict mean +/-SEM of at least two independent experiments (SH-SY5Y: n=4; KELLY: n=3; BE(2)-C: n=2).

To confirm the polyamine transport function of ATP13A3 in neuroblastoma cells, we assessed the effect of ATP13A3 overexpression on mediating the cytotoxic effects of high polyamine levels. Despite being essential for cell physiology and survival, at high intracellular concentrations, polyamines are cytotoxic ^18^. As an indirect measurement of polyamine uptake, we observed that upon polyamine supplementation, WT ATP13A3 overexpression, but not DN ATP13A3 overexpression increased the sensitivity of SH-SY5Y cells to increasing doses of putrescine and spermidine, while a similar trend was observed for spermine albeit not statistically significant **(Figure 3C – E**). Since we implemented heat-inactivated serum to neutralize extracellular polyamine oxidases (avoiding the extracellular conversion of polyamines into toxic hydrogen peroxide and aldehydes), the enhanced cytotoxic effects of polyamines in ATP13A3 WT versus DN expressing cells can be attributed to a higher intracellular polyamine content as a result of ATP13A3-mediated polyamine uptake. Indeed, similar findings were obtained when experiments were performed in the presence of 1 mM aminoguanidine, an inhibitor of polyamine oxidases, confirming that extracellular polyamine conversion is not responsible for the observed cytotoxicity **(SF 8)**.

To further validate the role of ATP13A3 in polyamine uptake in neuroblastoma cells, we performed transient siRNA-mediated ATP13A3 knockdown in *MYCN*-amplified (KELLY and BE(2)C) and non-*MYCN* amplified (SH-SY5Y) neuroblastoma cell lines **(SF 9)**. Compared with cells transfected with scrambled control siRNA (scr-ctrl), ATP13A3 knockdown cells had a significantly reduced uptake of radiolabeled putrescine (50 to 75% reduction) and especially spermidine (80 to 90% reduction) **(Figure 3F)**, which was also corroborated by analyzing uptake of BODIPY-labelled polyamines **(SF 10)**. Similar effects were observed in miRNA-mediated ATP13A3 stable knockdown cell lines **(SF 11)**.

Combined, our overexpression and silencing experiments provide strong evidence for an important role of ATP13A3 in basal polyamine uptake in neuroblastoma cells.

### 3.4 ATP13A3 supports neuroblastoma cell growth independent of MYCN

The finding that ATP13A3 expression in neuroblastoma patients is, independently of *MYCN* amplification, associated with worse survival suggests that ATP13A3-mediated polyamine uptake may play a part in neuroblastoma. To investigate this, we assessed the impact of ATP13A3 silencing on the growth and colony forming capacity of neuroblastoma cells. We observed that ATP13A3 silencing in both *MYCN*-amplified (KELLY) and non-*MYCN*-amplified (SH-SY5Y) neuroblastoma cells significantly attenuated cell growth and colony formation **(Figure 4A)**. These data suggest that targeting polyamine uptake alone impacts neuroblastoma viability and/or growth and that affected cells are not able to fully compensate for inhibition of polyamine uptake by upregulating polyamine biosynthesis. To provide further evidence for this, we assessed whether the polyamine uptake inhibitor AMXT 1501 ^33^ ^16^ ^34^ exerts similar effects on neuroblastoma cells. Indeed, AMXT 1501 as a single agent negatively impacted colony formation and viability of *MYCN*-amplified and non-*MYCN*-amplified neuroblastoma cells **(Figure 4B – C)**.

**Fig 4:**
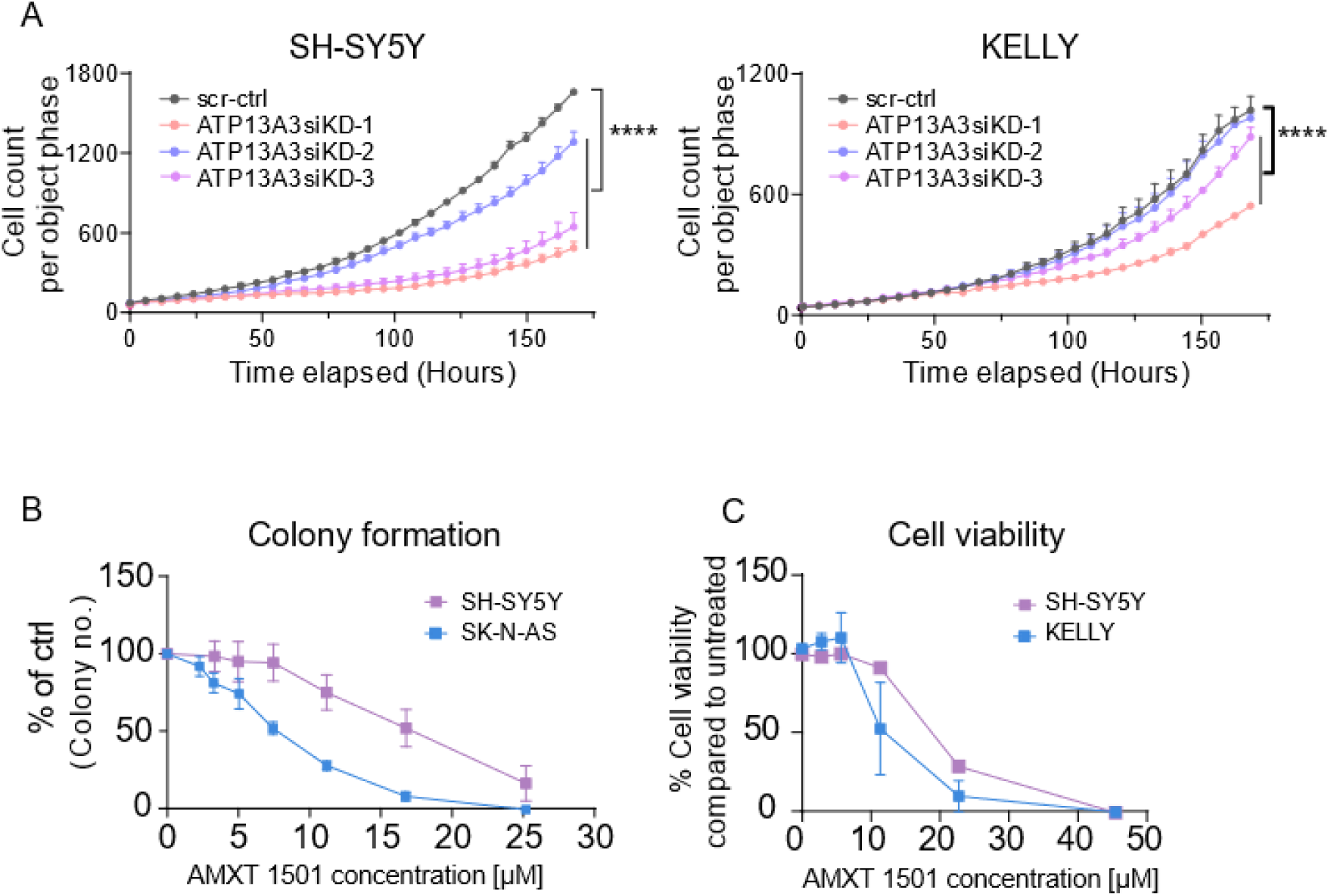
ATP13A3-mediated polyamine uptake supports the growth of *MYCN*-amplified and non-*MYCN*-amplified neuroblastoma cells. **(A)** Silencing of ATP13A3 decreases SH-SY5Y and KELLY cell growth, as assessed by the IncuCyte live cell imaging system. Two-way ANOVA was used for statistical analysis. Graphs depict mean +/-SEM of three independent biological replicates. **(B)** AMXT 1501 treatment reduced neuroblastoma cell colony formation ability in two non *MYCN*-amplified neuroblastoma cell lines (*MYCN*-amplified cell results published in Gamble et al paper ^15^). Graphs depict mean +/-SEM of three independent biological replicates. **(C)** AMXT 1501 treatment (72 hrs) reduced cell viability of *MYCN*-amplified (KELLY) and non-*MYCN*-amplified (SH-SY5Y) neuroblastoma cell lines, as assessed by cytotoxicity assay. Graphs depict mean +/-SEM of three independent biological replicates.

Hence, our data indicate that neuroblastoma cells depend on ATP13A3-mediated polyamine uptake to sustain their growth, implying that polyamine biosynthesis alone is not sufficient to provide these metabolites necessary for robust growth.

### 3.5 ATP13A3 mediates DFMO-induced polyamine uptake in neuroblastoma cells

Since the identity of the transporter(s) responsible for DFMO-induced polyamine uptake remain(s) unknown, we next evaluated the consequences of silencing ATP13A3 on DFMO-induced polyamine uptake. Knockdown of ATP13A3 alone was sufficient to prevent any DFMO-induced compensatory increase in radiolabeled putrescine and spermidine uptake in both *MYCN*-amplified and non-*MYCN* amplified neuroblastoma cells **(Figure 5A – D, SF12)**. Combined silencing of both ATP13A3 and SLC3A2 did not result in any further attenuation of uptake **(Figure 5A - D)**. These results indicate that ATP13A3 is at least partially responsible for the DFMO-induced polyamine uptake. In further support of this finding, ATP13A3 silencing also sensitized *MYCN*-amplified (KELLY) and non-*MYCN*-amplified (SH-SY5Y) neuroblastoma cells to escalating DFMO concentrations, which resulted in reduced growth and colony formation capabilities **(Figure 5E – F)**.

**Fig 5:**
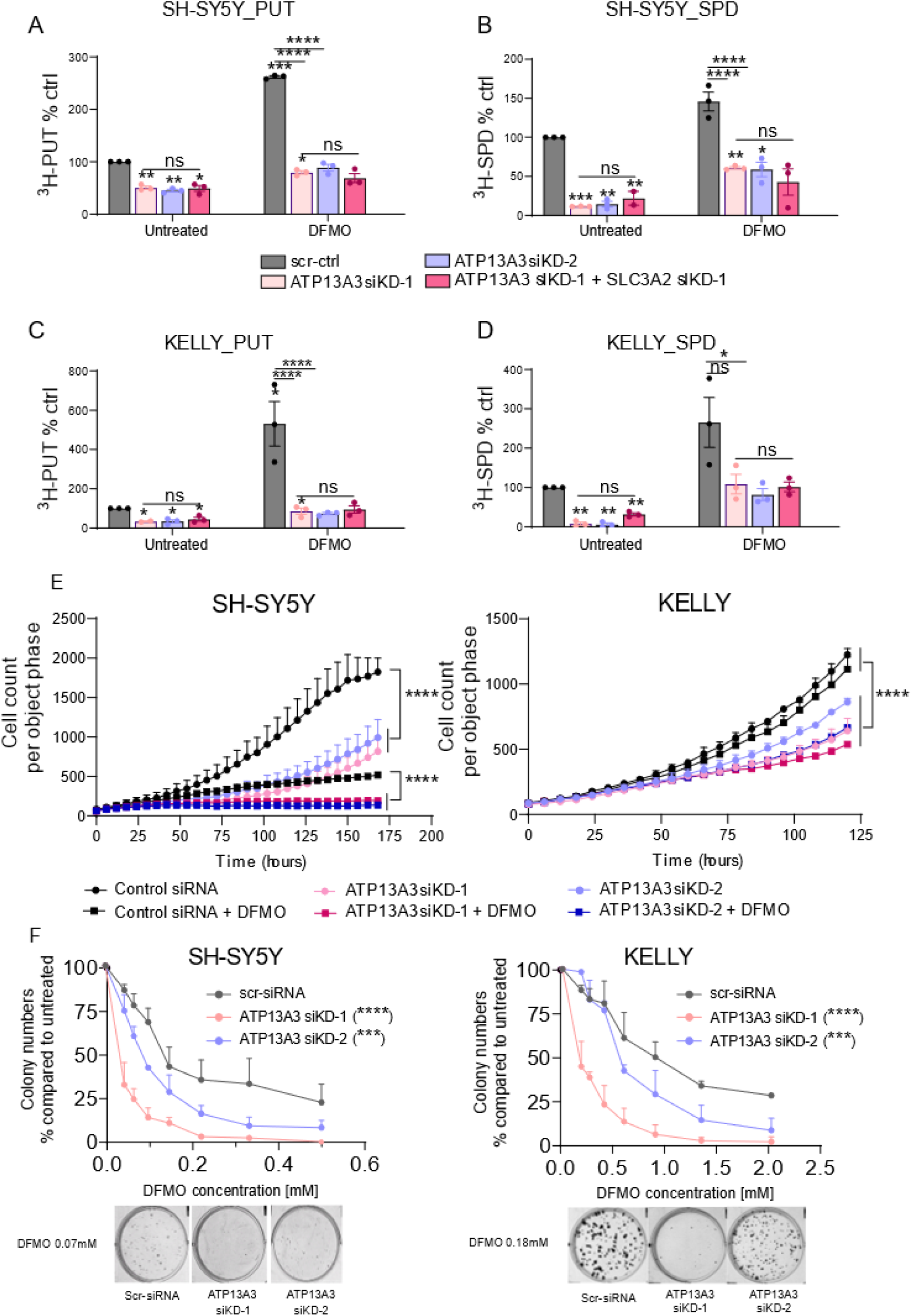
ATP13A3-mediated polyamine uptake contributes to DFMO-induced compensatory polyamine uptake in neuroblastoma cells and ATP13A3 silencing increases neuroblastoma cell sensitivity to DFMO. **(A – D)** siRNA-mediated silencing of ATP13A3 abrogates DFMO-boosted radiolabeled putrescine (PUT) and spermidine (SPD) uptake in SH-SY5Y and KELLY. Dual silencing of ATP13A3 and SLC3A2 does not decrease uptake levels further than ATP13A3 silencing does. One sample t-tests were used to compare uptake efficiency relative to untreated cells transfected with scr-ctrl (asterisks on the bar) One-way ANOVA was used for other group comparisons (asterisks between bars). Graphs depict mean +/-SEM of three independent biological replicates. **(E)** Silencing of ATP13A3 reduced cell growth upon treatment with DFMO as determined with the IncuCyte live cell imaging system. Two-way ANOVA was used to compare growth curves. Graphs depict mean +/-SEM of three independent biological replicates for SH-SY5Y and two independent biological replicates for KELLY. **(F)** ATP13A3 KD sensitizes neuroblastoma cells to DFMO treatment as measured by colony formation assays. Two-way ANOVA was used to compare dose-response curves between groups. Graphs depict mean +/-SEM of three independent replicates for SH-SY5Y, two independent experiments for KELLY.

Combined, our data demonstrate that ATP13A3 silencing counters the DFMO-induced upregulation of polyamine uptake, and enhances the efficacy of DFMO, suggesting that ATP13A3 might be a clinically relevant therapeutic target to potentiate polyamine biosynthesis inhibition approaches in neuroblastoma.

### 3.6 The polyamine transport inhibitor AMXT 1501 inhibits ATP13A3-mediated polyamine uptake in neuroblastoma cells

The polyamine transport inhibitor AMXT 1501 blocks DFMO-induced polyamine uptake ^16^ ^17^ and is currently being tested in combination with DFMO in clinical trials for solid cancers. Here we demonstrated that AMXT 1501 as a single agent inhibits neuroblastoma cell growth and colony formation similar to ATP13A3 silencing (**Figure 4B – C)**. Since the exact target(s) of AMXT 1501 remain(s) unknown and ATP13A3 emerges as a likely target, we next examined whether AMXT 1501 inhibits ATP13A3-mediated polyamine uptake. In line with our previous results and similar to ATP13A3 silencing ^15^ ^17^, AMXT 1501 effectively blocks putrescine and spermidine uptake and prevents the DFMO-induced compensatory increase in polyamine uptake in both *MYCN*-amplified and non-*MYCN* amplified neuroblastoma cells (**SF 13**). Reminiscent of ATP13A3 silencing, a sublethal dose of AMXT 1501 also sensitized *MYCN*-amplified and non-*MYCN*-amplified neuroblastoma cells to DFMO in colony formation assays demonstrating strong synergy between DFMO and AMXT 1501 in reducing neuroblastoma cell viability (**Figure 6A**).

**Figure 6:**
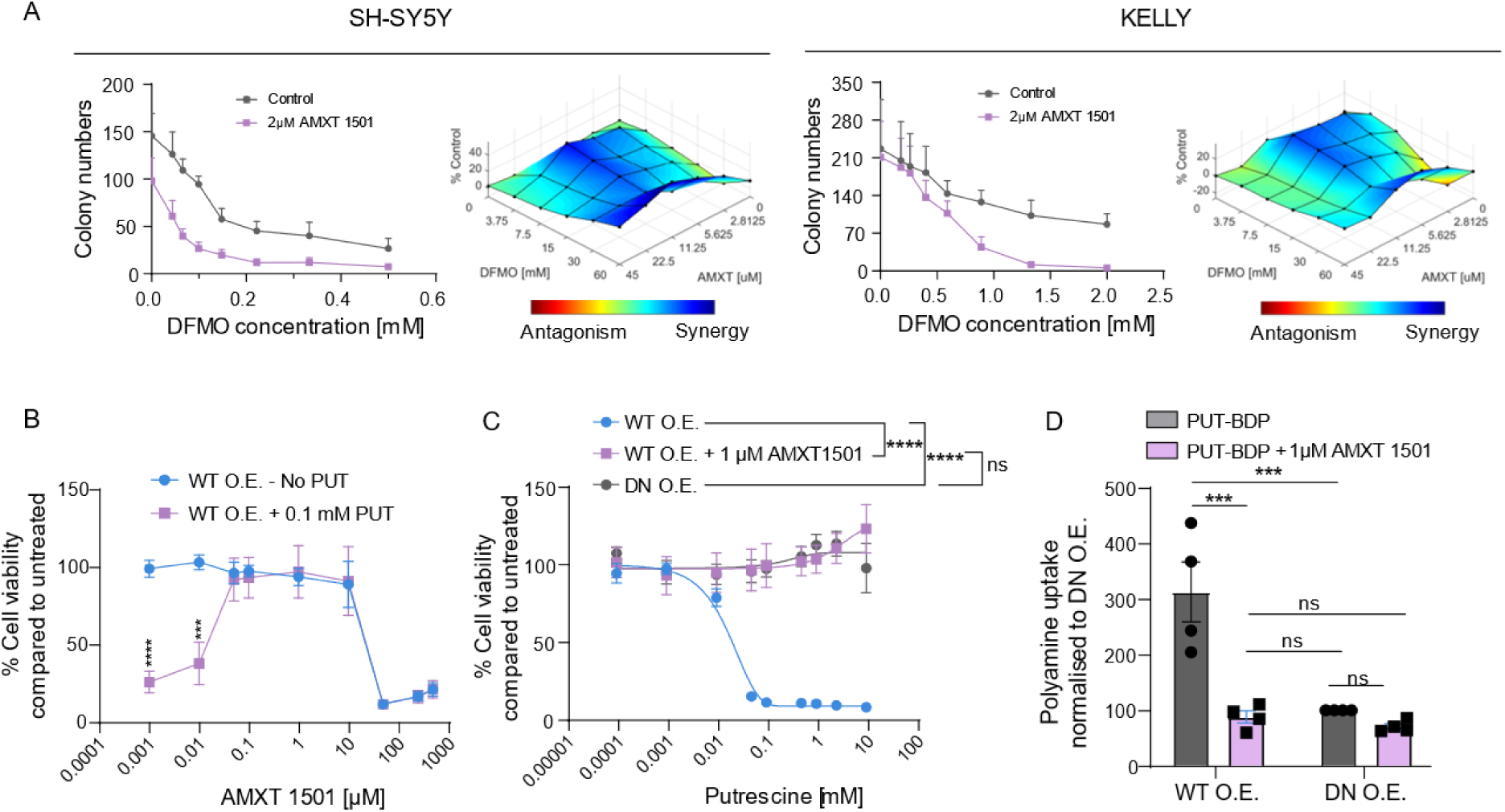
AMXT 1501 inhibits ATP13A3-mediated polyamine uptake. **(A Left**) A fixed dose of AMXT 1501 enhanced the inhibitory effect of DFMO on neuroblastoma colony formation. Graphs depict mean of three independent biological replicates. **(A Right)** DFMO and AMXT 1501 synergistically reduce neuroblastoma cell viability as assessed in 6x6 matrix resazurin-reduction based viability assays. Synergy is scored based on the Bliss Independency model. Graphs depict mean of three independent biological replicates **(B)** Cytotoxicity assay, using the MUH reagent to assess cell viability, showing the window of efficacy for AMXT 1501. Graphs depict mean +/- SEM of three independent biological replicates. **(C)** Overnight pre-treatment of SH-SY5Y cells overexpressing WT ATP13A3 with 1 µM AMXT 1501 protects them from toxic concentrations of putrescine (PUT). Two-way ANOVA was used to compare dose response curves in A and B. mean +/-SEM of three independent biological replicates. **(D)** Overnight pre-treatment with 1 µM AMXT 1501 abolishes ATP13A3-mediated PUT-BDP. One sample t-tests were used to compare mean uptake levels relative to untreated DN O.E. cells. One-way ANOVA was used to compare mean uptake levels for all other multiple comparisons. Graphs depict mean +/-SEM of four independent biological replicates.

To provide more conclusive evidence for AMXT 1501 inhibiting ATP13A3-mediated polyamine uptake in neuroblastoma, we next assessed whether the activity of ATP13A3 can be effectively hindered by AMXT 1501. We determined the uptake inhibition capacity of AMXT 1501 by evaluating the potential of AMXT 1501 to abrogate the cytotoxic effects of putrescine when supplemented at a concentration normally lethal for cells overexpressing ATP13A3 **(Figure 6B)**. We defined the concentration of AMXT 1501 required to prevent 50% of putrescine toxicity by blocking ATP13A3-mediated polyamine import, *i.e.* the IC_50_ for polyamine transport inhibition, to be approximately 0.05 μM. Additionally, AMXT 1501 by itself exerted cytotoxic activity with IC_50_ value around 50 μM. Based on these findings, we proceeded to pre-treat ATP13A3 overexpressing SH-SY5Y cells with a non-toxic concentration of 1 μM AMXT 1501 that is able to block polyamine transport of putrescine. Interestingly, we completely rescued the putrescine toxicity in ATP13A3 overexpressing SH-SY5Y cells to the point that they were indistinguishable from ATP13A3 DN overexpressing cells **(Figure 6C)**. Similarly, AMXT 1501 treatment completely inhibited ATP13A3-mediated BODIPY-labelled putrescine uptake **(Figure 6D).**

Our data thus provide compelling evidence that AMXT 1501 effectively targets ATP13A3-mediated polyamine uptake, underscoring the therapeutic promise of ATP13A3 targeting in neuroblastoma.

In conclusion, this study identified ATP13A3 as a critical regulator of basal and DFMO-induced polyamine uptake in *MYCN*-amplified and non-*MYCN*-amplified neuroblastoma cells, which is targetable by the polyamine transport inhibitor AMXT 1501, and thereby highlights ATP13A3 as a novel therapeutic target for neuroblastoma.

## 4. Discussion

### ATP13A3 is a major polyamine transporter in neuroblastoma that can be blocked by AMXT 1501

Neuroblastoma cells are addicted to polyamines for their growth and survival and these cells respond to inhibition of polyamine biosynthesis upon treatment with DFMO by upregulating polyamine uptake as a rescue mechanism to prevent polyamine depletion ^26^ ^27^. In this study we compared the role of candidate polyamine transporters SLC3A2 and ATP13A3 in neuroblastoma polyamine uptake under basal and DFMO-treated conditions. Although high expression of *SLC3A2* is associated with poorer outcome in neuroblastoma patients and SLC3A2 knockdown reduced polyamine uptake in neuroblastoma cells, SLC3A2 silencing did not abolish the DFMO-induced increase in polyamine uptake. In contrast to SLC3A2, silencing of ATP13A3 was sufficient to prevent the compensatory increase in polyamine uptake under DFMO treatment. Also, overexpression of ATP13A3 increased intracellular polyamine levels and enhanced polyamine uptake in basal conditions, whereas ATP13A3 knockdown resulted in reduced polyamine uptake. Our study therefore highlights the critical role of ATP13A3 in polyamine uptake in neuroblastoma cells under basal and DFMO-treatment conditions, pointing to ATP13A3 as a primary polyamine transporter responsible for polyamine uptake in neuroblastoma. Importantly, ATP13A3-mediated polyamine uptake can be prevented by AMXT 1501, a polyamine uptake inhibitor currently in phase I/II clinical trials for cancer, indicating that preventing polyamine uptake via ATP13A3 may at least partially explain the mode of action of the drug. Based on our findings, ATP13A3 emerges as a major polyamine transporter in neuroblastoma that is a target of the drug AMXT 1501.

### ATP13A3 emerges as a new therapeutic target for neuroblastoma and other cancers

ATP13A3 knockdown reduces the growth and colony formation capacity of neuroblastoma cells, which is consistent with high ATP13A3 expression being predictive of poor outcome in neuroblastoma patients. Also, the polyamine transport inhibitor AMXT 1501, which acts at least in part via ATP13A3, inhibits neuroblastoma cell growth and colony formation showing that polyamine uptake in basal conditions is important for neuroblastoma growth. Additionally, ATP13A3 knockdown increased the sensitivity of neuroblastoma cells to DFMO pointing to synergistic effects of blocking polyamine uptake via ATP13A3 and inhibiting polyamine synthesis with DFMO. These results indicate that inhibition of ATP13A3, possibly in combination with DFMO, may be of therapeutic interest for neuroblastoma.

Studies in other cancers including pancreatic, head, neck and cervical cancers, have noted associations between ATP13A3 expression and poorer overall survival, suggesting a role for this polyamine uptake pathway in other cancer types as well, although detailed studies into the role of this transporter in most cancers are largely lacking ^35^ ^36^ ^20^ ^37^ ^38^. ATP13A3 was highlighted as the major importer of spermidine and spermine in human pancreatic cells, and metastatic pancreatic cancer cells displayed higher polyamine import and expression of ATP13A3 as compared to cells with slow proliferation. ATP13A3 knockdown significantly decreased the growth of human pancreatic cancer cells under DFMO treatment ^38^. Also, other members of the P5B-ATPase family have been linked to different cancers ^6^ ^21^. ATP13A2 has been implicated in melanoma, colon cancer, hepatocellular carcinoma, acute myeloid leukemia and non-small-cell lung cancer ^39^ ^40^ ^41^ ^42^ ^43^. We also reported that high ATP13A4 expression explains the increased polyamine uptake in the breast cancer cell line MCF7 compared to the non-tumorigenic epithelial breast cell line MCF10A ^21^. ATP13A4 amplification has been frequently observed in patients with non-small-cell lung cancer ovarian cancer, cervical cancer, head and neck cancer, endometrial cancer, uterine endometrioid carcinoma, bladder cancer, melanoma, and breast cancer ^21^.

### Towards P5B-type ATPase inhibitors for cancer therapy

The discovery of ATP13A3 as the primary driver in the hyperactive polyamine transport system of neuroblastoma may pave the way for developing more targeted strategies to attack cancer cells that heavily rely on increased polyamine import. To deprive neuroblastoma cells of their polyamine addiction, selective ATP13A3 inhibitors can be developed. Although the safety of AMXT 1501 (in combination with DFMO) has been established in phase I/II clinical trials (NCT03536728; NCT05500508), targeting ATP13A3 with specific inhibitors may not be without risk, since *ATP13A3* gene mutations have been associated with the pathogenicity of pulmonary arterial hypertension ^44^. Future analysis of conditional *Atp13a3* knockout mice may help establish the (patho)physiological consequences of long term versus short term *Atp13a3* deficiency.

Inhibition of ATP13A3 could be achieved either with polyamine analogues that may occupy the polyamine binding site or with unrelated small molecules targeting catalytic sites of the transporter. Another potential strategy that harnesses the role of ATP13A3 in neuroblastoma involves linking polyamines with toxic warheads to direct a treatment specifically to cells highly dependent on polyamine import ^45^ ^46^. Crucially, the distinct polyamine preferences and cancer-type specific expression profiles exhibited by different P5B-ATPases can potentially be harnessed to design effective polyamine analogues. In cellular polyamine uptake experiments, subtle differences between ATP13A2, ATP13A3 and ATP13A4 have been observed in terms of their polyamine specificity. ATP13A2 exhibits the highest preference for spermine and spermidine, ATP13A4 enables uptake of putrescine, spermidine and spermine, while based on both previously published data and our own findings in ATP13A3 overexpressing SH-SY5Y cells, ATP13A3 appears to mainly facilitate putrescine and spermidine uptake ^18^ ^19^ ^23^. However, Sekhar et al. observed that ATP13A3 displays the least preference for putrescine, behaving more similarly to ATP13A2 with a preference for spermine and spermidine ^20^. The precise polyamine specificity of ATP13A3 therefore remains uncertain and might even depend on the cellular context of the transporter or co-expression with other transporters. Determining the differences in polyamine specificity will require conducting biochemical assays on purified ATP13A3 protein, which could also pave the way towards inhibitor screening and for confirming AMXT 1501’s direct impact on ATP13A3.

### ATP13A3 as a potential biomarker or therapeutic target under DFMO treatment

DFMO is currently under investigation in a clinical trial in children with neuroblastoma in combination with standard of care chemotherapy and dinutuximab (NCT05500508). Studies by us and others unequivocally demonstrated that cancer cells in general, and neuroblastoma cells in particular, compensate for inhibition of polyamine synthesis by boosting uptake of polyamines from the extracellular environment ^26^ ^27^. The compensatory polyamine uptake phenomenon can be explained by the tight regulation of intracellular polyamine levels by the interplay between antizyme and antizyme-inhibitor. Levels of antizyme, an enzyme that inhibits both polyamine synthesis and polyamine uptake, are regulated by intracellular polyamine levels ^47^. This mechanism is mediated by the enzyme called Antizyme Inhibitor, the expression of which increases after polyamine depletion upon DFMO treatment, resulting in increased polyamine uptake ^48^ ^49^ ^50^. Here, we show a role for ATP13A3 in the DFMO-induced polyamine uptake in neuroblastoma cells suggesting that ATP13A3 may contribute to the resistance mechanisms employed by neuroblastoma cells following DFMO treatment. Therefore, combination therapies with DFMO and ATP13A3 inhibitors may be of therapeutic interest to prevent this compensatory response. Moreover, information regarding ATP13A3 expression or activity may help delineate which patients are more likely to develop rescue mechanisms to polyamine biosynthesis inhibition treatments with DFMO.

While it remains unclear whether ATP13A3 levels can be used to predict responsiveness to DFMO, we noted a trend towards an increase in ATP13A3 protein levels in response to DFMO treatment of neuroblastoma cells (data not shown). In pancreatic cancer cells however, the expression level of ATP13A3 correlated positively with the sensitivity to polyamine transport inhibitors. To inhibit growth, pancreatic cancer cells with higher ATP13A3 expression required higher concentrations of polyamine transport inhibitors but lower concentrations of DFMO ^38^. Conversely, in this study in neuroblastoma, we observed in neuroblastoma cells that ATP13A3 knockdown increases sensitivity to DFMO, suggesting that high ATP13A3 expression might drive resistance to DFMO. These contradictory findings in neuroblastoma versus pancreatic cancer cells highlight the importance to thoroughly assess the functions of polyamine transporters in different cancer types.

To further validate ATP13A3 as a therapeutic target and assess its biomarker value in predicting responsiveness to DFMO in neuroblastoma, *in vivo* xenograft studies will be required to determine the efficacy of DFMO, AMXT 1501 and their combination in a panel of patient-derived xenograft models with varying expression levels of ATP13A3 and/or ODC1.

### ATP13A3 is a target of the *MYC* oncogene

The mechanisms regulating ATP13A3 expression are poorly understood. Our analysis of ChIP-seq databases of *MYCN*-amplified and c-MYC-expressing neuroblastoma cells, suggests that both MYCN and c-MYC can directly bind the promotor of *ATP13A3* and thus may regulate *ATP13A3* mRNA. Supporting this finding, *ATP13A3* mRNA levels were significantly higher in *MYCN*-amplified versus non-*MYCN*-amplified patients. The MYCN-polyamine axis is of particular significance in neuroblastoma research, since multiple genes within the polyamine pathway, including *ODC1* and *SLC3A2*, are direct targets of MYCN ^32^. The link between MYCN and the polyamine pathway is so well established that targeting the polyamine homeostasis pathway is seen as an indirect method of targeting MYCN which has been deemed undruggable ^51^ ^52^. In relation to ATP13A3 however, high *ATP13A3* expression levels negatively correlate with survival of neuroblastoma patients, irrespective of *MYCN* status. Moreover, multivariate analysis showed that elevated *ATP13A3* expression is an independent predictor of poor prognosis in neuroblastoma, independent of *MYCN* amplification. Also, ATP13A3 knockdown appears effective in both *MYCN*-amplified and non-*MYCN* amplified neuroblastoma cell lines. While our ChIP-seq database analyses point towards c-MYC/MYCN directly regulating ATP13A3 expression through binding to the gene’s promotor, dedicated promotor-binding studies as previously performed for other polyamine pathway genes are required to provide firm evidence for this ^15^.

In conclusion, our data show that ATP13A3 represents an interesting novel therapeutic target to inhibit the polyamine transport system in neuroblastoma to increase the efficacy of DFMO treatment and limit neuroblastoma growth.

## Author Contributions

M.A. and W.G. are shared first authors and M.H., P.V. and K.S. are shared last and co-corresponding authors. W.G., M.A., L.X., P.V. and K.S. conceptualized the study with contributions of R.P., M.D.N and M.H. The stable cell line models were designed and generated by C.V.d.H. S.V. synthesized provided the BODIPY-labelled polyamines. All experiments were performed by W.G. and M.A., with support by M.K., A.K., D.S., E.R. and A.B. X.G. and C.M. performed multivariate survival and ChIPseq database analyses. W.G. and M.A. analyzed the data. M.A. and W.G. wrote the first draft of the manuscript, which has been revised and edited by P.V. and K.S. All authors have read and approved the manuscript.

## Supplementary Figures

**Supplementary Figure 1:**
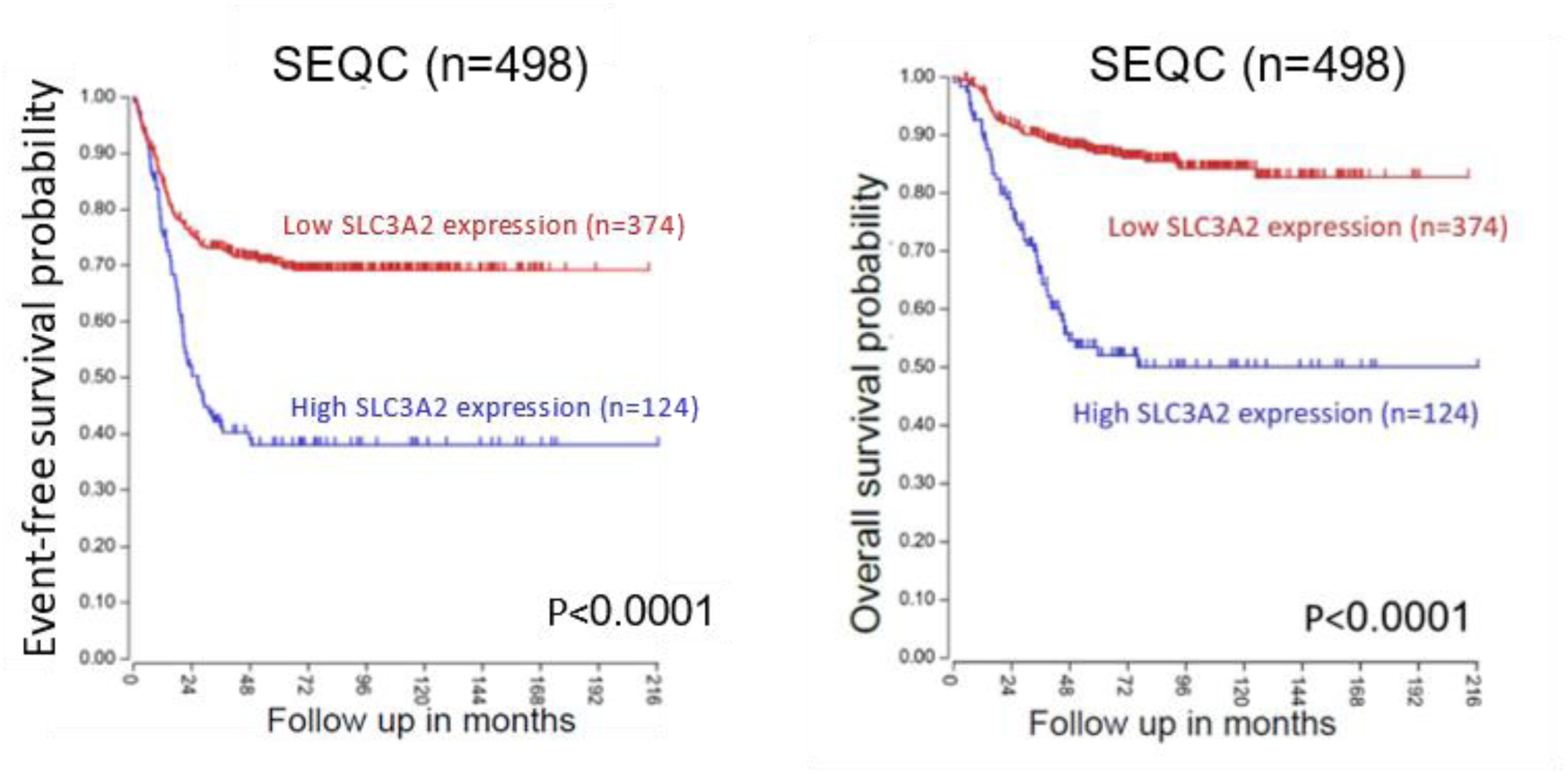
High *SLC3A2* expression is associated with worse overall and event-free survival in neuroblastoma patients (SEQC dataset n=498). Patients were dichotomized around the upper quartile (UQ) of gene expression. Log-rank test was used to compare survival curves.

**Supplementary Figure 2:**
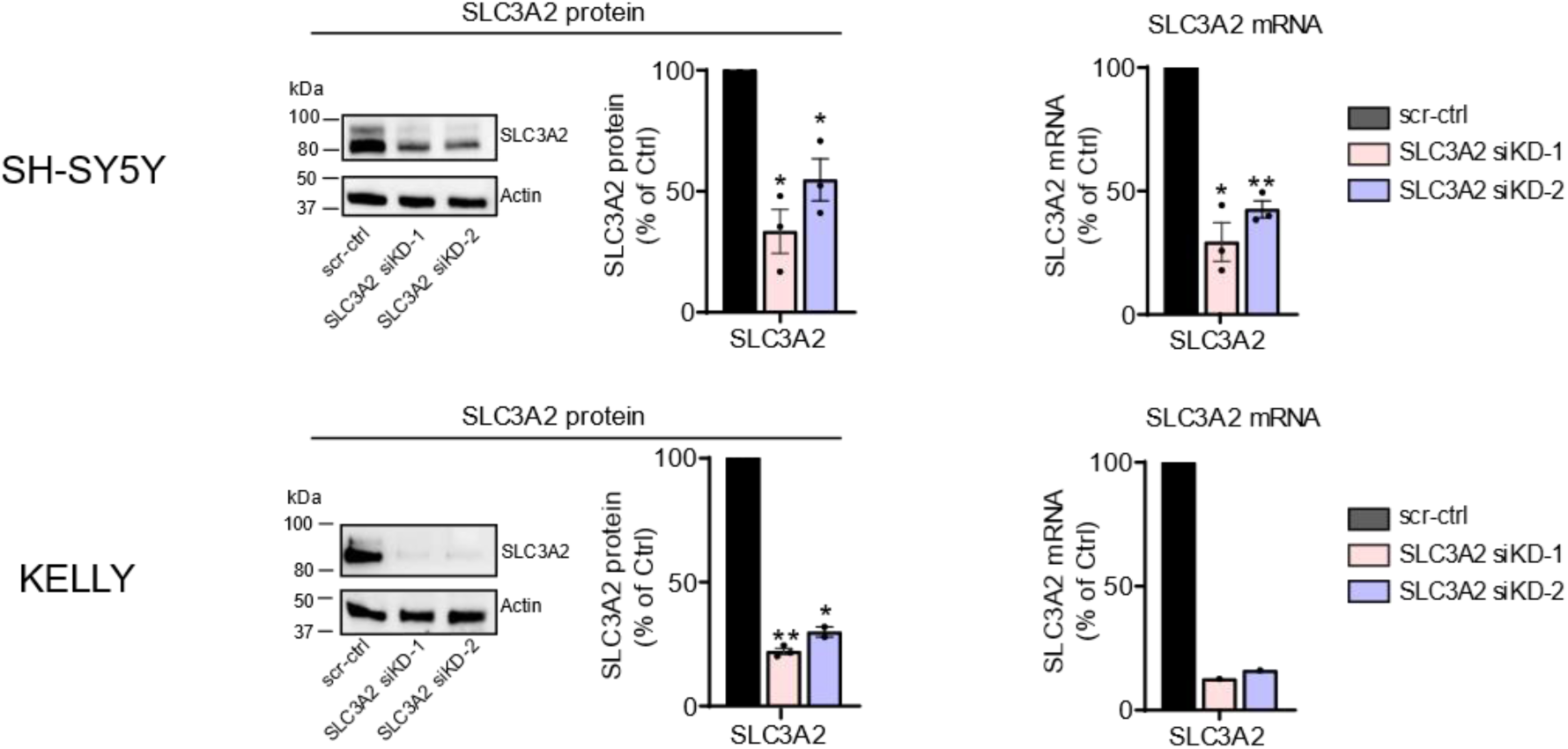
Validation of effective SLC3A2 knockdown in SH-SY5Y and KELLY cells. Reduced SLC3A2 protein (left) and mRNA (right) levels after siRNA-mediated silencing (for 48 hr) in SH-SY5Y (top) and KELLY (bottom) cells. One sample t-test was used to assess the significance of silencing relative to scr-ctrl transduced cells. Presented immunoblots are representative of results obtained in three independently performed experiments. Graphs depict mean +/-SEM of n=3 independent biological repeats except for the measurement of SLC3A2 mRNA levels in KELLY (n=1).

**Supplementary Figure 3:**
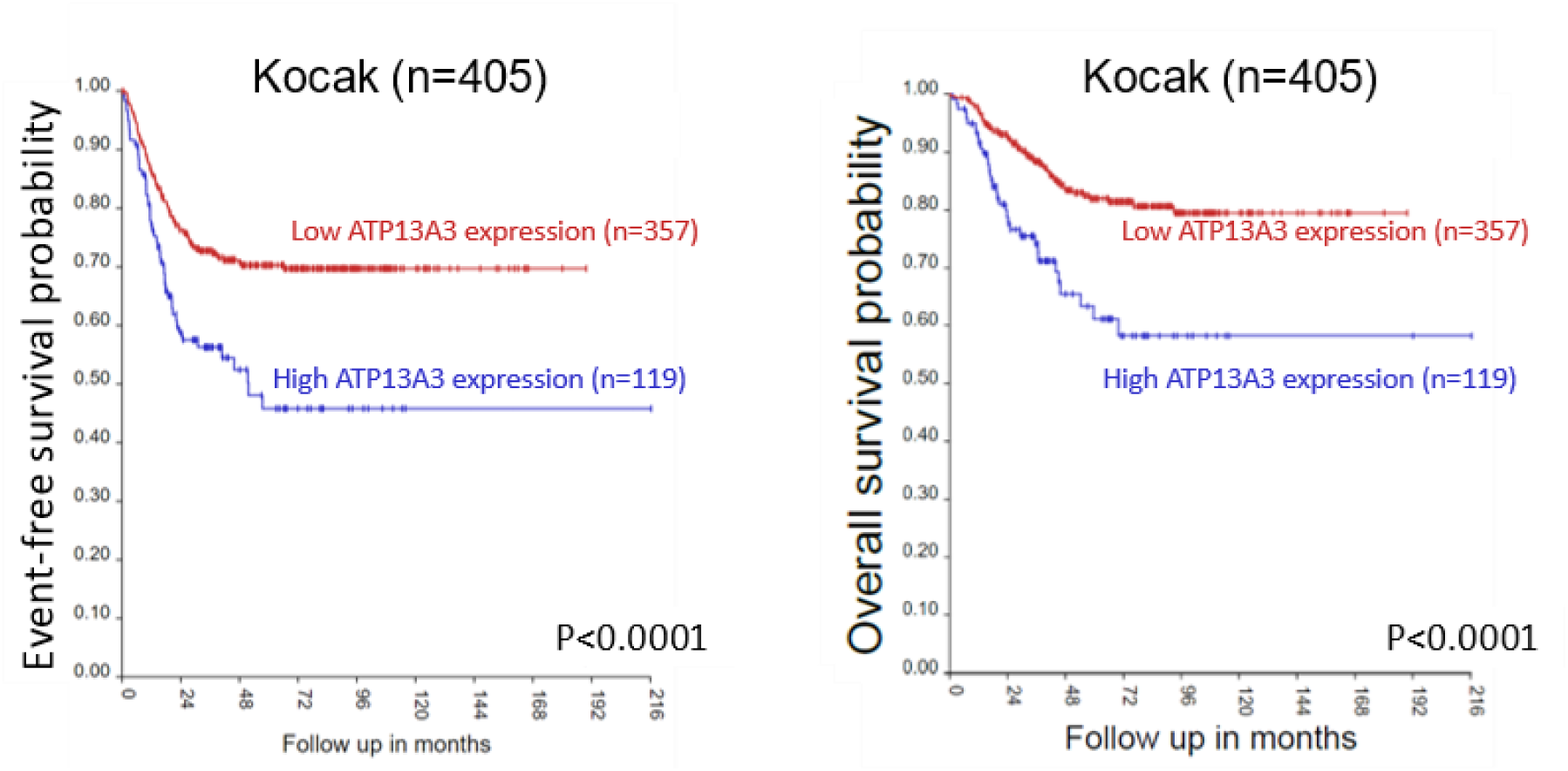
High expression of *ATP13A3* is associated with significantly worse outcome in neuroblastoma patients. **(A – B)** High expression level of *ATP13A3* is associated with worse event-free and overall survival in the Kocak (n=405, B) patient cohorts. Patients were dichotomized around the upper quartile (UQ). Log-rank test was used to compare survival curves.

**Supplementary Figure 4:**
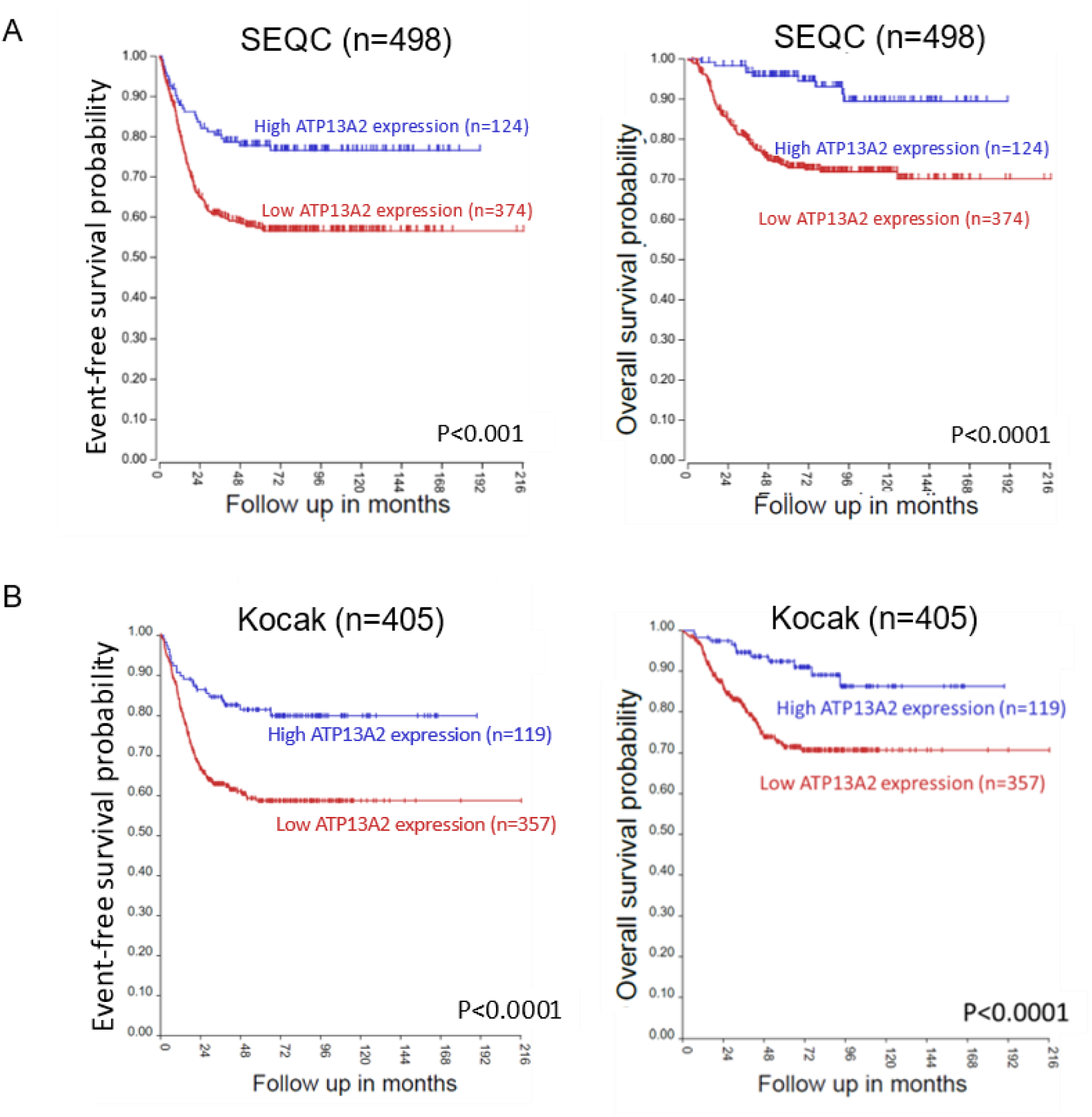
High expression of *ATP13A2* is not associated with a worse prognosis in neuroblastoma patients. Survival curves for high and low *ATP13A2* expression in the **(A)** SEQC (n=498) and **(B)** Kocak (n=649) cohorts. Log-rank test was used to compare survival curves.

**Supplementary Figure 5:**
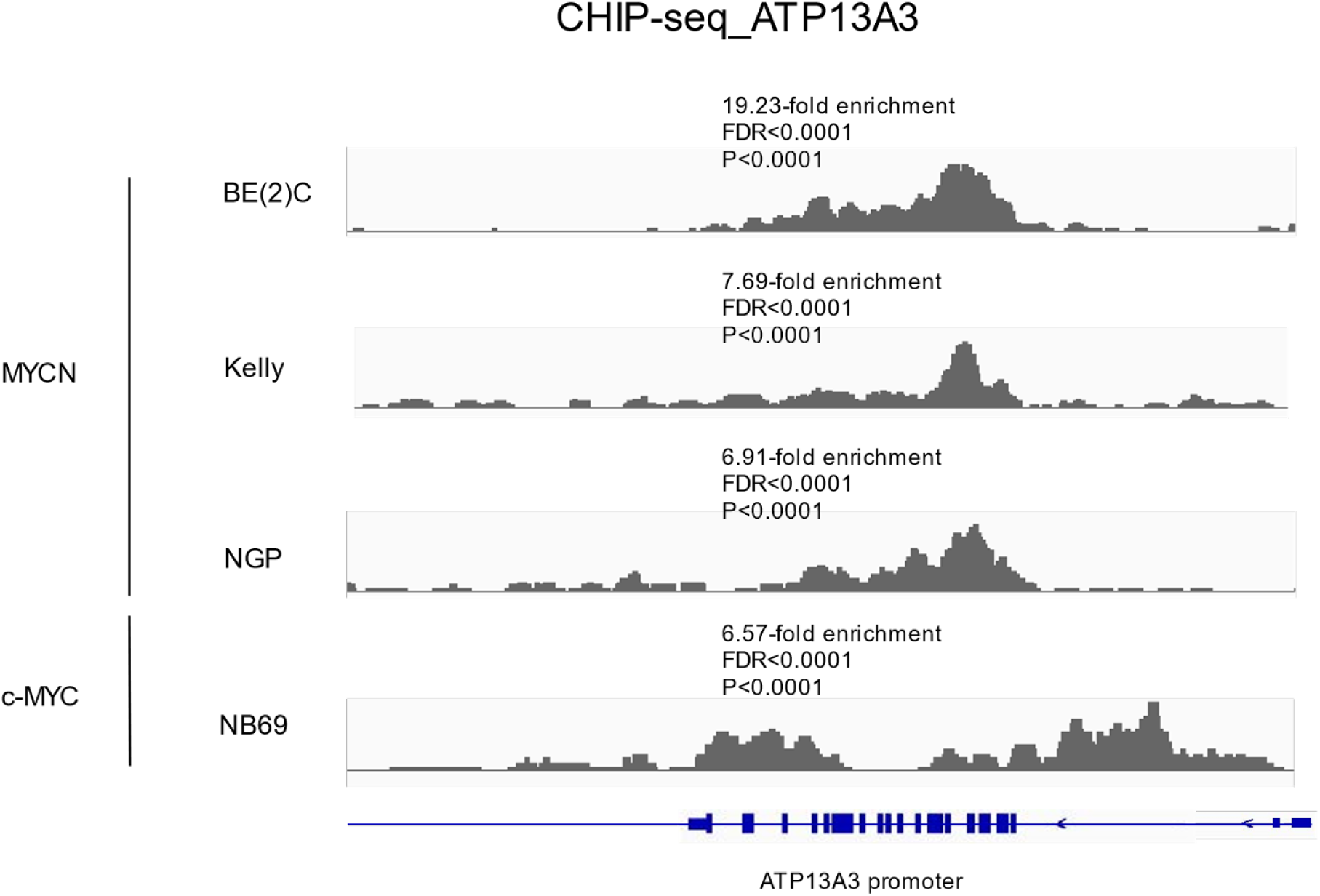
MYCN and cMYC bind the *ATP13A3* promotor in *MYCN*-amplified and cMYC-expressing neuroblastoma cells. ChIP-seq tracks of ATP13A3 after pulldown with anti-MYCN in three *MYCN*-amplified neuroblastoma cell lines, BE(2)-C, KELLY, and NGP, and with anti-c-MYC antibody in NB69 neuroblastoma cells. Phred quality scores with a threshold of q30 were used to decrease the chance of false positives.

**Supplementary Figure 6:**
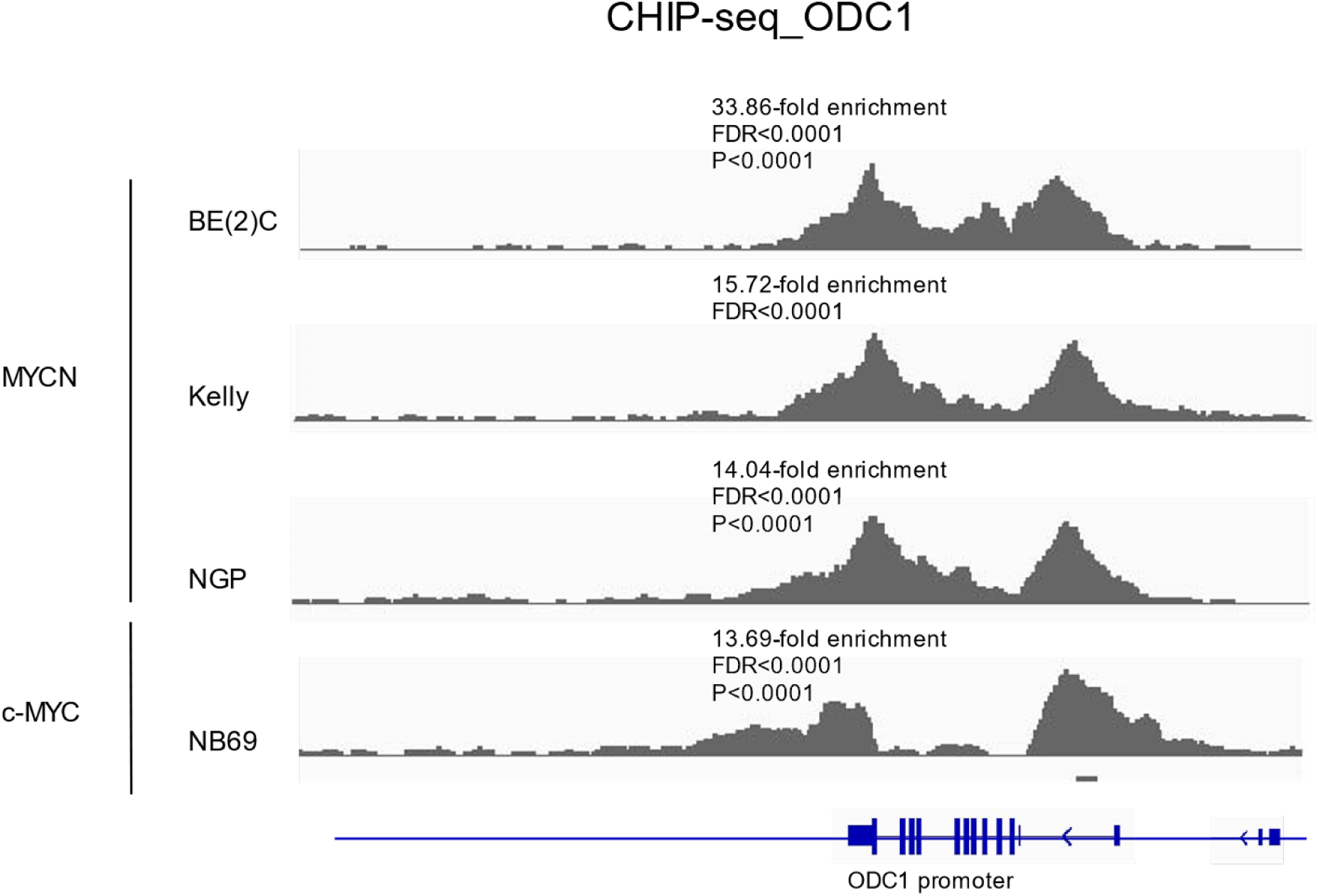
MYCN and cMYC bind the ODC1 promotor in neuroblastoma cells. ChIP-seq tracks of ODC1 after pulldown with anti-MYCN in three *MYCN*-amplified neuroblastoma cell lines, BE(2)-C, KELLY, and NGP, and with anti-c-MYC antibody in NB69 neuroblastoma cells. Phred quality scores with a threshold of q30 were used to eliminate false positives.

**Supplementary Figure 7:**
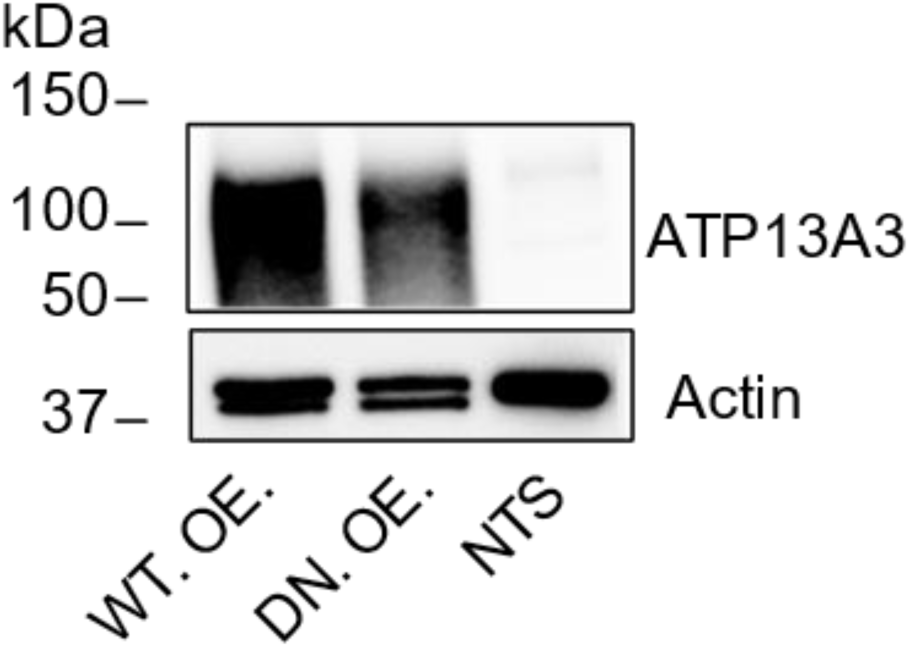
Validation of SH-SY5Y cells overexpressing WT ATP13A3 and the catalytically dead (DN) mutant of ATP13A3. Western Blot showing ATP13A3 expression in the overexpression models vs. the nontransduced (NTS) control.

**Supplementary Figure 8:**
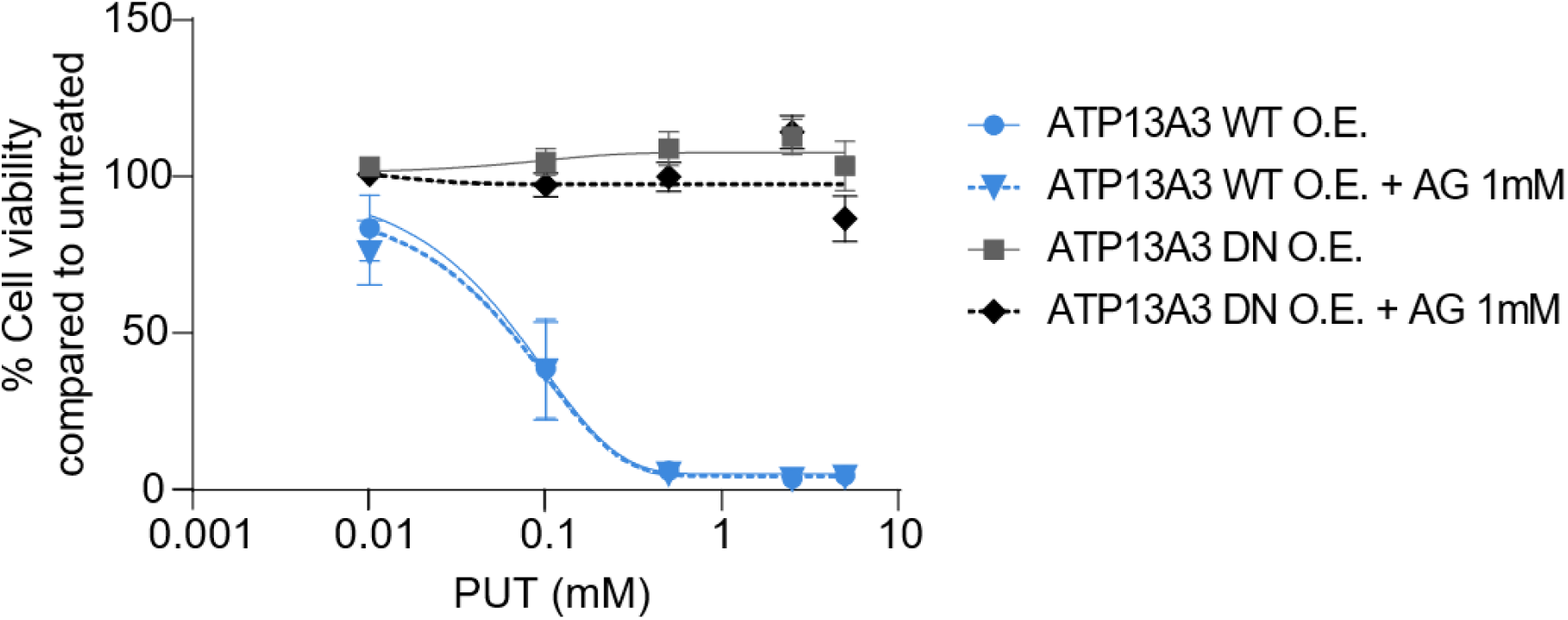
Increased sensitivity to polyamine toxicity upon WT ATP13A3 overexpression is not due to extracellular polyamine oxidases. No significant difference was seen in PUT toxicity for either of the two WT or DN overexpressing (O.E.) SH-SY5Y cells with or without 1 mM aminoguanidine supplementation. Graphs depict mean +/-SEM of n=3 independent biological repeats.

**Supplementary Figure 9:**
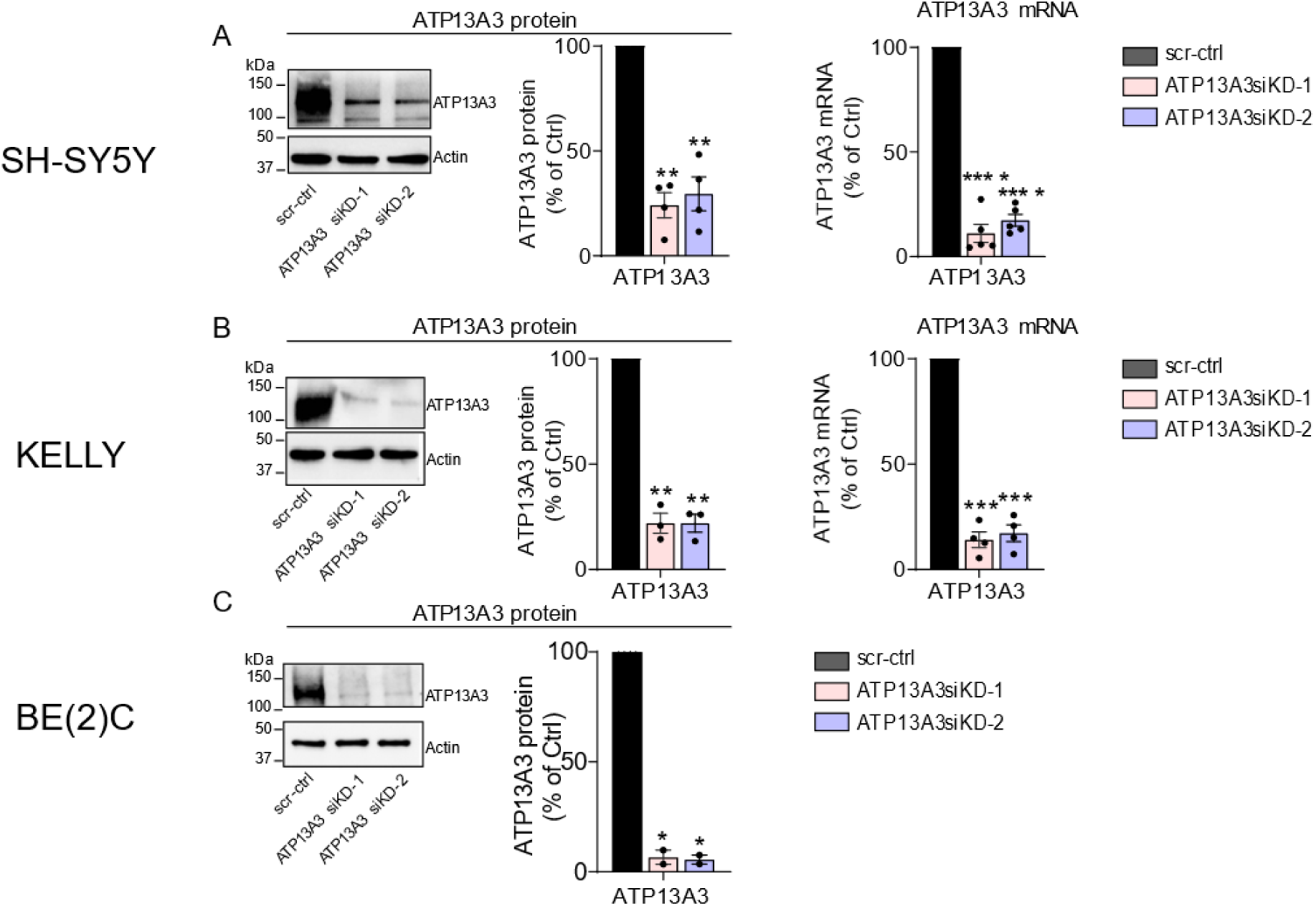
Validation of siRNA-mediated ATP13A3 knockdown. Reduced ATP13A3 protein (left) and mRNA (right) levels after siRNA-mediated silencing in neuroblastoma cells (48 hrs). One sample t-test was used to assess the significance of silencing relative to scr-ctrl transduced cells. The depicted blot is representative for data obtained in at least two independently performed experiments (SH-SY5Y: n=4; KELLY: n=3; BE(2)-C: n=2). ATP13A3 mRNA graphs depict mean +/- SEM of at least four independent biological replicates.

**Supplementary Figure 10:**
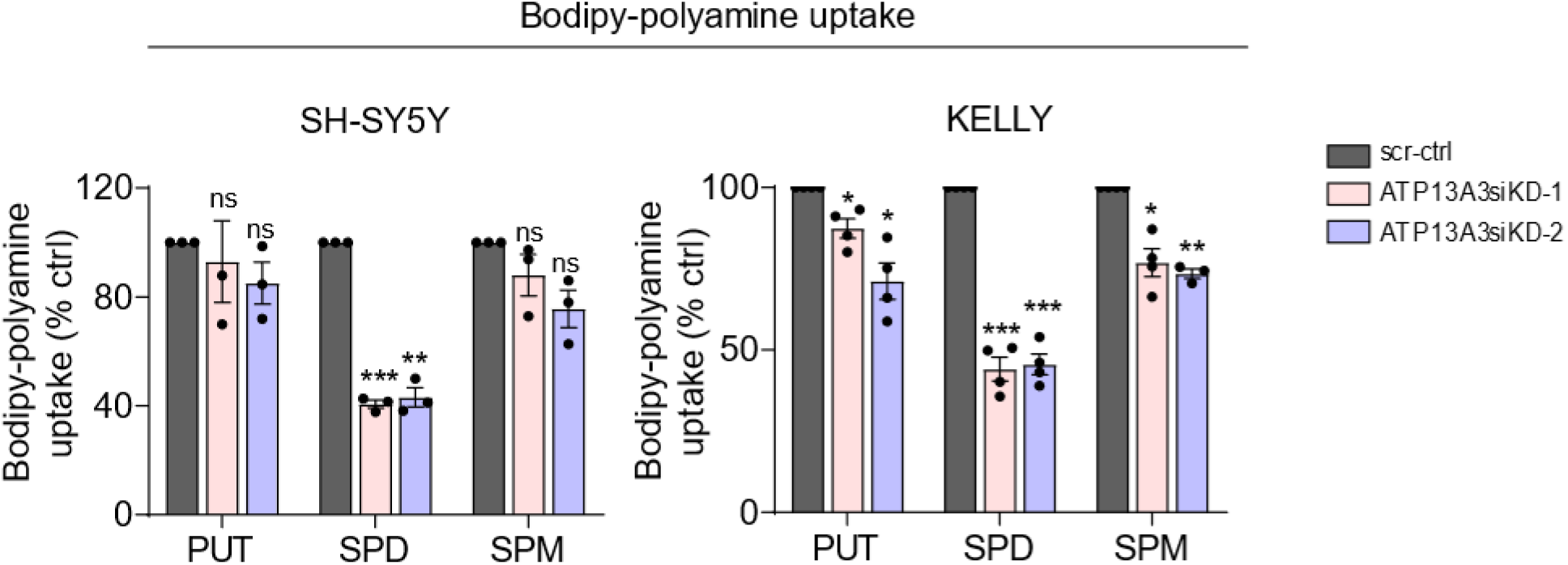
Transient siRNA-mediated ATP13A3 knockdown inhibits polyamine-BDP uptake in SH-SY5Y and KELLY cells. One sample t-test was used to establish the significance of the decrease in uptake upon ATP13A3 silencing relative to cells transduced with scr-ctrl. Graphs depict mean +/-SEM of at least three independent biological replicates.

**Supplementary Figure 11:**
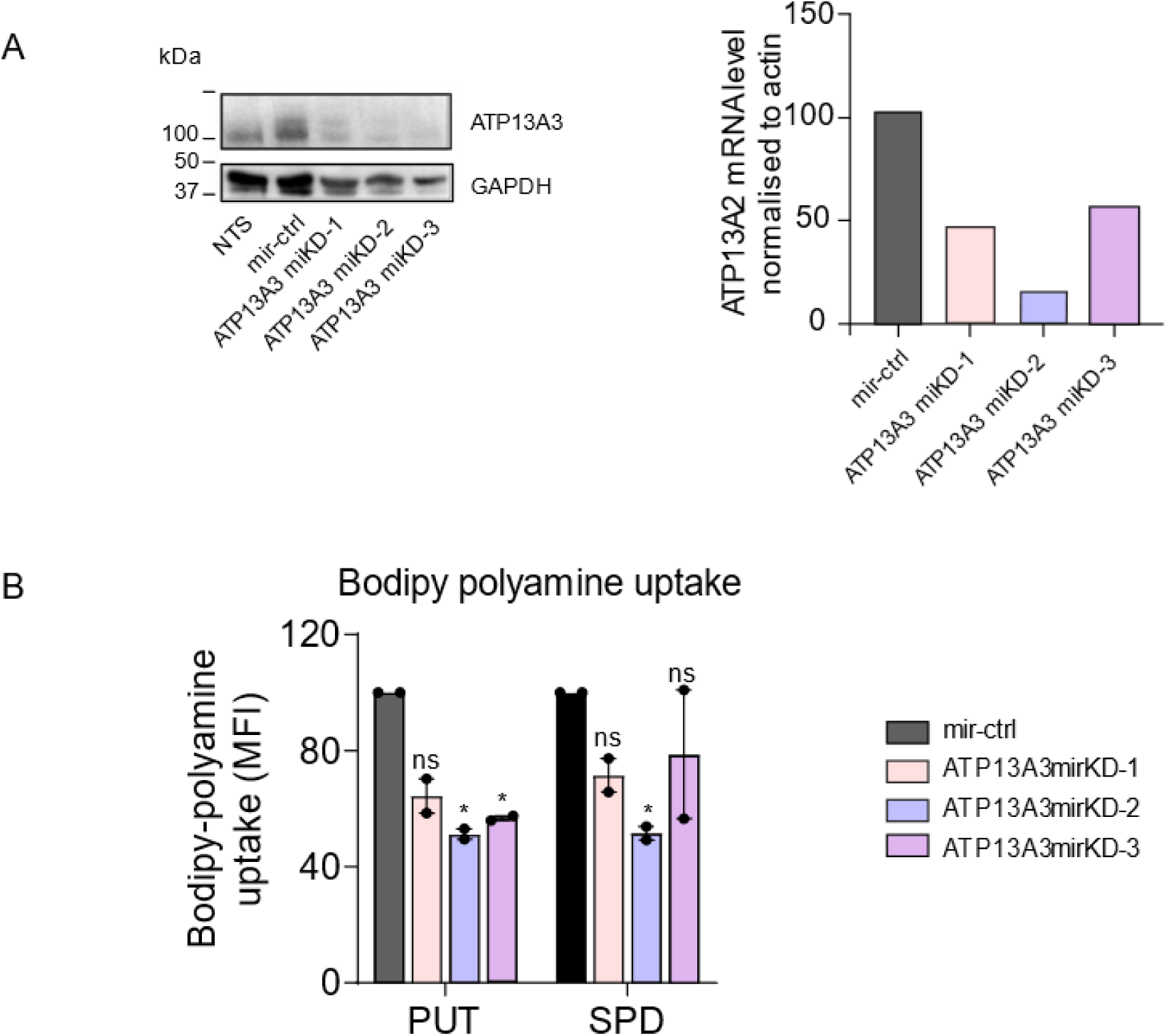
**(A)** Protein (left) and mRNA (right) levels of ATP13A3 following stable lentiviral transduction to knockdown ATP13A3 in SH-SY5Y cells. **(B)** Confirmation of reduced polyamine-BODIPY uptake in SH-SY5Y cells with stable miRNA-mediated ATP13A3 knockdowns. One sample t-test was used for statistical analysis. Graphs depict mean +/-SEM from two independent experiments.

**Supplementary Figure 12:**
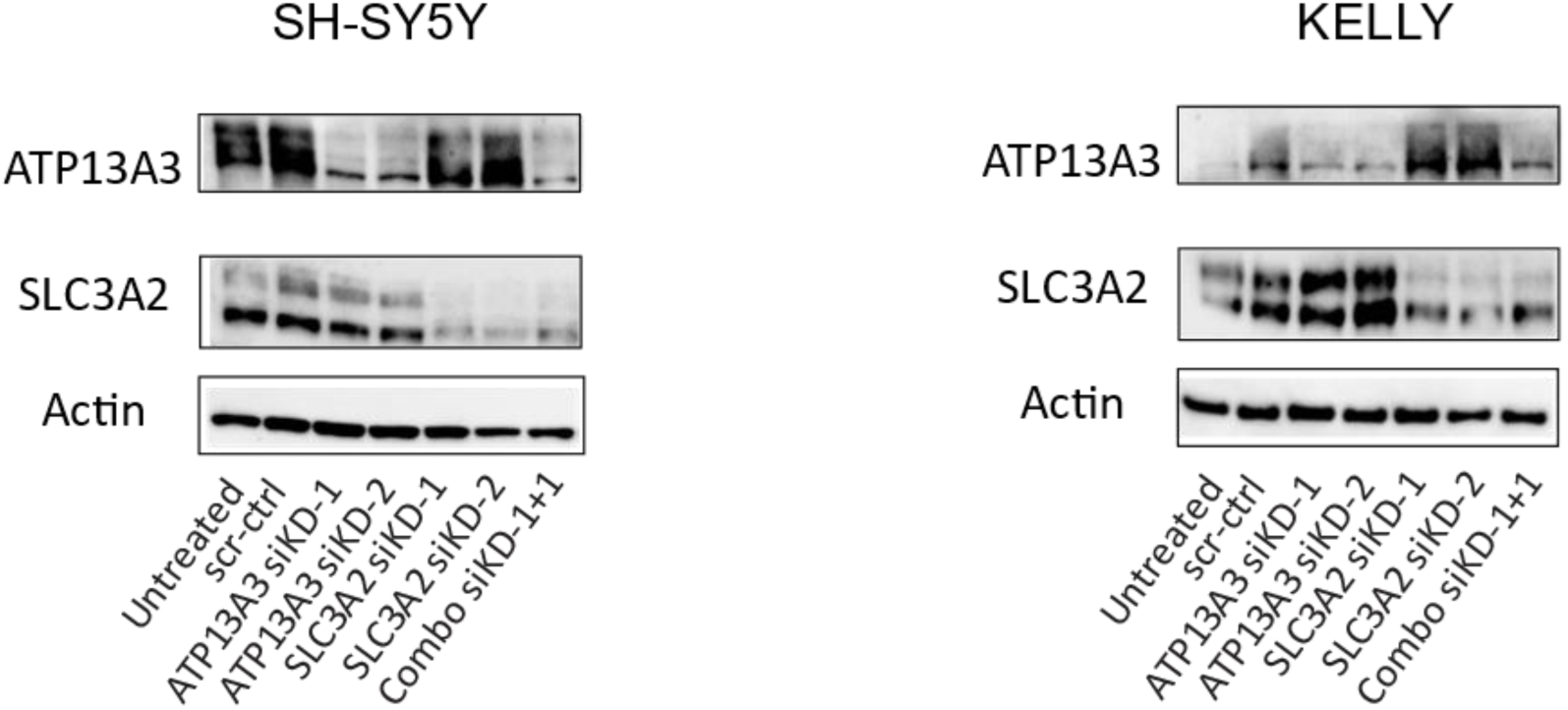
Immunoblotting confirming effective siRNA-mediated silencing of ATP13A3 and/or SLC3A2 in SH-SY5Y and KELLY cells. Immunoblot is representative for two independent experiments with protein expression evaluated 48hrs post transfection. Combo indicates dual ATP13A3 and SLC3A2 silencing.

**Supplementary Figure 13:**
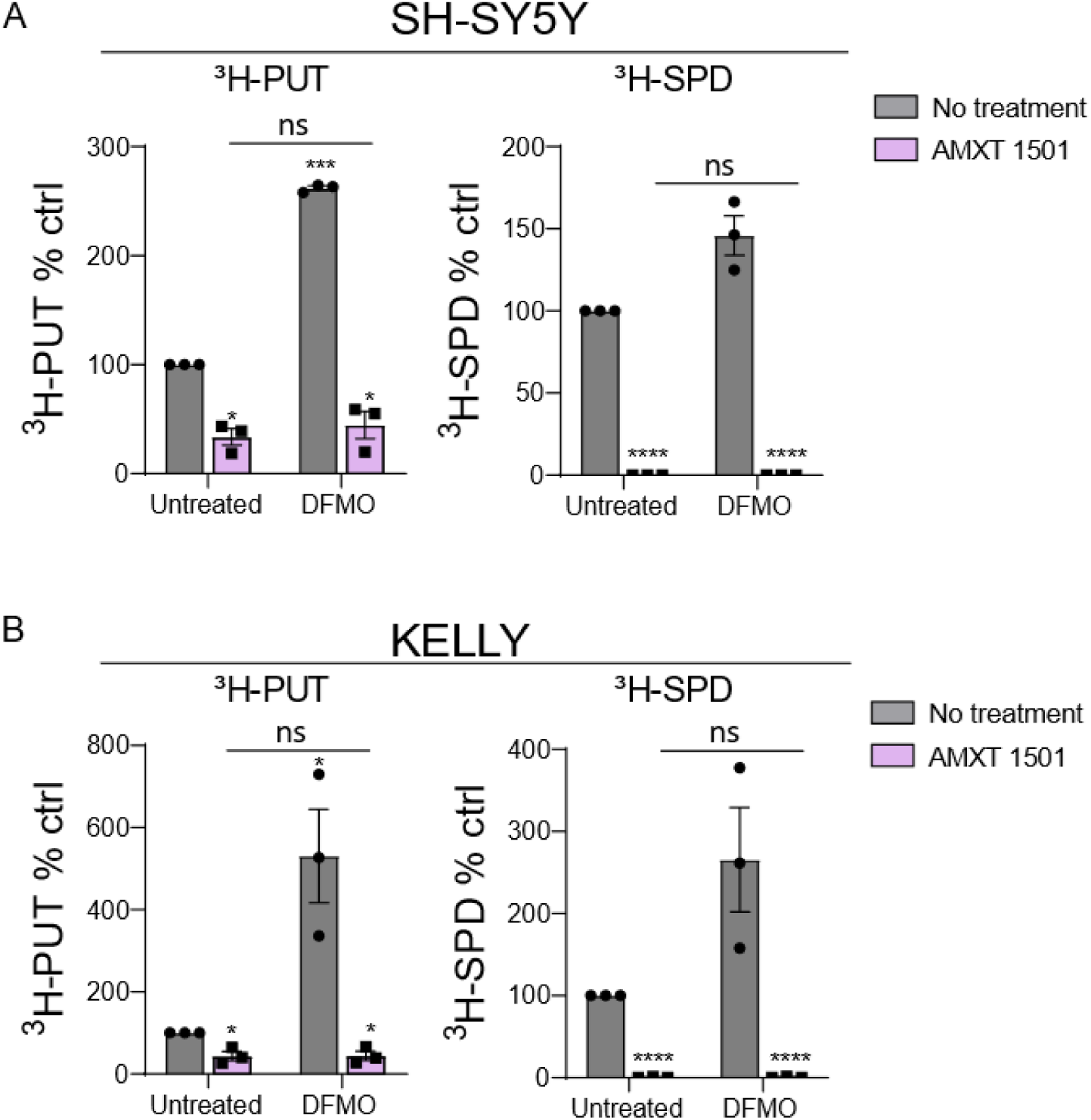
Overnight pre-treatment with 1 µM AMXT 1501 inhibits baseline and DFMO-induced polyamine uptake in **(A)** non-*MYCN*-amplified and **(B)** *MYCN*-amplified neuroblastoma cells. One sample t-test was used for comparing mean uptake levels relative to untreated cells and one-way ANOVA is used for other group comparisons. Graphs depict mean +/-SEM of three independent biological replicates.

